# Claustrum Morphometry Reveals Spatially Heterogeneous Alterations in 22q11DS

**DOI:** 10.64898/2026.10.27.762130

**Authors:** Aaron Althauser, Farnaz Delavari, Caren Latrèche, Dimitri Van De Ville, Stephan Eliez

**Affiliations:** Developmental Imaging and Psychopathology Laboratory, Department of Psychiatry, University of Geneva School of Medicine, Geneva, Switzerland; Medical Image Processing Laboratory, Neuro-X Institute, École Polytechnique Fédérale de Lausanne, Geneva, Switzerland; Radiology Department, Lausanne University Hospital and University of Lausanne, Lausanne, Switzerland; Department of Radiology and Medical Informatics, University of Geneva, Geneva, Switzerland; CIBM Center for Biomedical Imaging, Geneva, Switzerland; Department of Genetic Medicine and Development, University of Geneva School of Medicine, Geneva, Switzerland; Fondation Pôle Autisme, Geneva, Switzerland

**Keywords:** Claustrum, 22q11.2 Deletion Syndrome, Surface-Based Morphometry, Working Memory

## Abstract

The human claustrum is a small, irregular, and complexly positioned structure, yet it carries long range projections to most cortical regions responsible for executive functioning. Prior structural MRI research has largely relied on global volume as the metric of its morphology. Such measures may obscure regionally specific morphological changes relevant to developmental and pathological variation.

We therefore addressed this limitation by applying longitudinal surface-based morphometry to 874 scans from 440 individuals aged 5–35 years, comparing typical development with 22q11.2 deletion syndrome (22q11DS), a genetic condition characterized by brain dysmaturation, and assessing associations with working memory performance.

Our data show a significant age-related volumetric decline that is reduced in 22q11DS left claustra, while surface deformation and thickness analyses revealed spatial patterns unique to hemisphere, group, and developmental age. In controls, the claustrum surface deformed with age in clustered zones along a dorsoventral gradient, and the dorsal-anterior zone thinned across development while the ventral zone thickened. 22q11DS claustra showed less age-related deformation than controls, whose claustrum surface changes associated with verbal working memory. This structural correlate was statistically equivalent to zero in carriers, consistent with a disrupted claustral development to the working memory vulnerability of the syndrome.

These findings show the value of surface-based morphometry over volume alone, highlight dysmaturation and regionally organized claustrum shape variation, and suggest that disrupted structural organization is accompanied by a weaker claustrum-working memory association.

## 1 Introduction

The claustrum is a thin, irregular sheet-like structure located between the putamen medially and the insular cortex laterally. Its extensive connectivity has historically motivated the hypothesis that it mediates sensory integration and consciousness (Crick and Koch 2005; Liaw and Augustine 2023). Despite its small size, the claustrum exhibits a broad topographic organization, variable neurochemistry, and overlapping projection gradients (Dávila et al. 2005; Mathur et al. 2009; Erwin et al. 2021; Marriott et al. 2021; Grimstvedt et al. 2023; Fodoulian et al. 2025). In humans, dorsal-anterior zones project to frontal, prefrontal, motor, and premotor cortices (Fernández-Miranda et al. 2008), whereas ventral and posterior claustral zones show stronger projections to visual, auditory, and temporal processing areas in mice (Wang et al. 2017).

This topographic connectivity is also consistent with a broader hypothesized role of the claustrum as a consolidator of distributed cortical networks (Rodríguez-Vidal et al. 2024). Mouse studies have been important models for linking together the underlying cellular, circuit, and behavioral mechanisms. For example, specific claustral neurons and their inhibition have been related to working memory performance (Bhattacharjee et al. 2026), and top-down processing has been linked to the role of specific cell groups within the claustral complex that shape claustrum-frontal microcircuits (Atlan et al. 2018, 2024; Jackson et al. 2018; White et al. 2018; White and Mathur 2018). In humans, claustrum activity and connectivity have been implicated in salience processing (Madden et al. 2022), cognitive control, attention, and working memory (Krimmel et al. 2019); however, human neuroimaging studies of the claustrum are limited.

Further translational studies show that claustro-frontal circuits mature dynamically, with ACC input switching from excitation to inhibition as claustro-PV interneurons mature (Shaker et al. 2026), making the claustrum a target for age-sensitive abnormalities in executive function and cognitive control. Volumetric, connectivity, and neurochemical abnormalities have been reported in Parkinson’s and Alzheimer’s diseases, schizophrenia, autism spectrum disorder, and attention-deficit/hyperactivity disorder (ADHD) (Firbank et al. 2005; Cascella et al. 2011; Smythies et al. 2012; Wegiel et al. 2014; Bernstein et al. 2016; Wang et al. 2016; Arrigo et al. 2017; Chon et al. 2017; Kang & Sponheim 2017; Liu et al. 2017, 2021; Sitte et al. 2017; Atilgan et al. 2022; Ayyildiz et al. 2023; Kartal et al. 2024; Mazurek et al. 2025; Schinz et al. 2025; Shelley et al. 2025). As such, all prior *in vivo* human MRI studies have assessed the claustrum by global volume. However, the claustrum is not a homogeneous structure, so global volumetric measures may obscure spatially specific morphological changes.

A regionally resolved description of the claustrum surface is better suited to localize structural changes that volumetric analyses may average out. Moreover, interpreting these disease-related volume differences requires normative reference data spanning the same age range: a reduced or enlarged claustrum is meaningful only relative to its typical developmental or aging trajectory. Lifespan research indicates that claustrum volume follows distinct growth and decline trajectories, with sex differences and a normative rightward asymmetry (Ayyildiz et al. 2026). Mapping claustrum surface variation across development could reveal fine-grained morphometric markers of atypical maturation. Such markers could clarify how regional changes contribute to the cognitive and psychiatric impairment observed in at-risk populations.

Therefore, we aimed to study longitudinal changes in regional shape measures that quantify claustrum morphology and their relationship to typically developing individuals and patients with 22q11.2 deletion syndrome (22q11DS), a rare genetic disorder that confers a high risk of schizophrenia and specific deficits in executive functioning, working memory, and attention (Majerus et al. 2007; Maeder et al. 2016; Montojo et al. 2017). We further investigated how these surface-based measures correlated with differences in working memory task performance between healthy controls and 22q11DS carriers.

## 2 Materials and Methods

### 2.1 Participants

A total of 874 scans from genetically confirmed 22q11DS patients (N = 465; 190 subjects) and healthy controls (HC; N = 409; 250 subjects) were enrolled from an ongoing longitudinal study conducted in Geneva with follow-up scans every three years. Participant ages ranged from 3–59 years, restricted to 5–35 for the age-matched analytical sample. Controls comprised unaffected siblings and a community sample recruited through the Geneva state school system, matched to 22q11DS on age and sex, and screened to exclude personal history of psychiatric disorder, developmental delay, or congenital cardiac anomaly.

### 2.2 Cognitive Measures

Global intellectual functioning was estimated using age- and date-appropriate Wechsler intelligence scales (Wechsler 2019). Digit Span, administered as part of the same scales, yielded two subscores: Digit Span Forward (DSF), repeating digits in presented order, and Digit Span Backward (DSB), repeating them in reverse. DSF indexes short-term phonological storage, while DSB additionally requires active mental manipulation and indexes verbal working memory processes (Baddeley 1996). A Total Digit Span score (DST) summed DSF and DSB. Spatial Span Total (SST) scores were computed from participants’ performance on a task which indexes visuo-spatial working memory, recorded using the same Wechsler scales on a subset of adults.

### 2.3 MRI Acquisition

Structural MRI data were acquired across the 19-year follow-up using three 3T Siemens platforms: Trio (N = 290) and Prisma Fit (N = 256) at the Hôpitaux Universitaires de Genève (HUG); and Prisma (N = 249) at the Fondation Campus Biotech Geneva. All images were acquired using a T1-weighted MPRAGE sequence: voxel size = 0.9 x 0.9 x 1.1 mm3, TR = 2500 ms, TE = 3 ms, flip angle = 8 deg, slices = 192, slice thickness = 1.1 mm, FOV = 220 mm, phase-encoding direction R > L. The Trio-era acquisitions (2007-2014) used the same vendor MPRAGE sequence.

### 2.4 Quality Control

An initial pool of 965 sessions with clinical metadata entered the analysis pipeline. Sessions were excluded in stages. First, 45 sessions failed Euler quality control (Euler Z < -1.5; Rosen et al. 2018). Second, 13 subjects were excluded because of registration failure or bad-quality visits (mean residual > 1.5 mm), which removed all their sessions from template construction (N = 27), as the analysis pipeline required successful mesh registration and within-subject template convergence. Finally, age-matching the subject pool to ages 5–35 removed 19 sessions. This yielded 874 sessions from 440 participants with acceptable quality requirements.

### 2.5 Segmentation, Vertex-Alignment, and Template Construction

Data preprocessing from acquisition to analysis is detailed in Fig. 1. We used a dedicated claustrum segmentation model (Mauri et al. 2025) available in Freesurfer v8.0 to generate per-session segmentations across the full dataset; segmentations were manually inspected by a radiologist trainee (FD), topologically corrected, and converted to triangular surface meshes using FreeSurfer’s Marching Cubes implementation (Montani et al.1994).

**Fig. 1.**
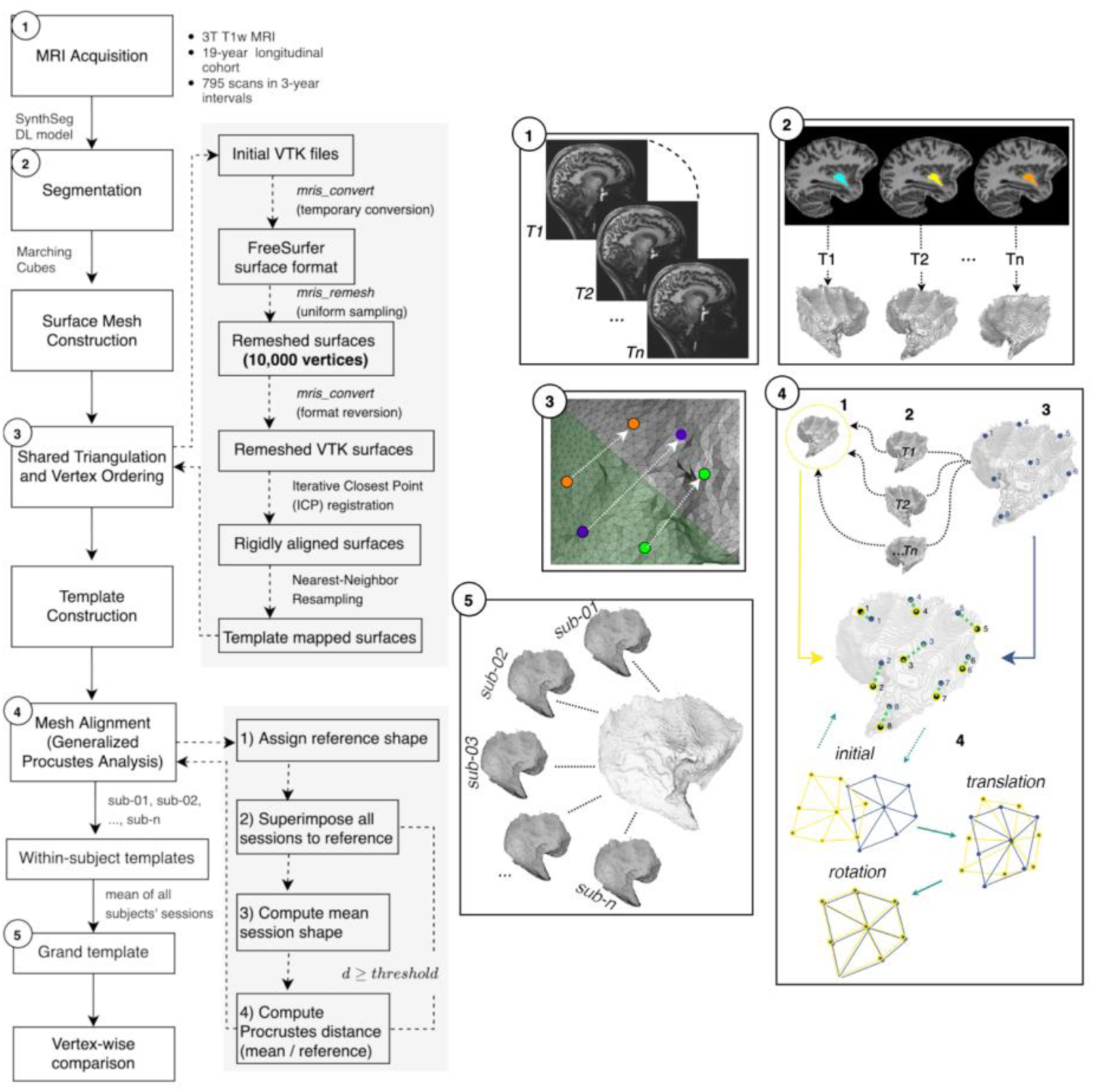
Overview of the Method. The diagram shows how we processed data from initial acquisition to vertex-wise comparison via template construction.

To account for multiple sessions, we adapted a mesh-based analog of FreeSurfer’s unbiased longitudinal template (Reuter et al. 2012) using Iterative Closest Point (ICP) registration, constructed subject-level templates via generalized Procrustes analysis (Gower 1975), and constructed a grand template from the mean of all within-subject templates. For downstream comparisons, we used the HC template as a normative reference; statistical outputs were unchanged under a group-balanced template. Scanner distortion from multi-site acquisition was appropriately assessed and did not bias the template construction (Suppl. Mats. Section 1.1).

### 2.6 Surface-Based Measures

At each of the 10,000 points on the left and right claustrum surfaces, we computed the amount of variation between a subject and the reference, yielding a signed value per point. Negative values indicate inward (perpendicular) deformation of the subject’s surface relative to the reference template, and positive values indicate outward deformation away from the mean reference (Fig. 2b). At each vertex, the displacement was obtained by:

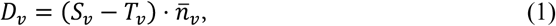

**Fig. 2.**
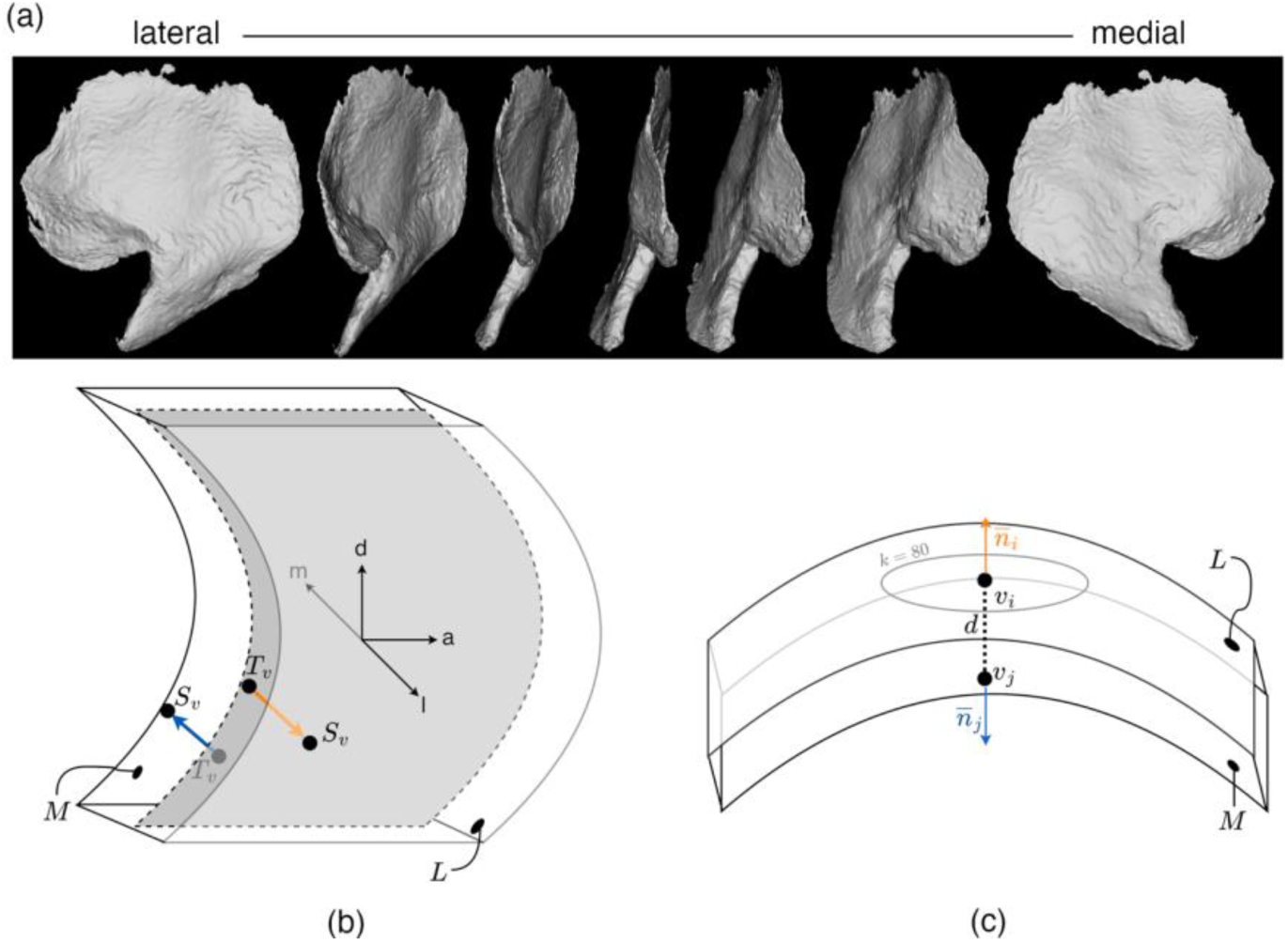
Surface geometry. (a) The grand template 3D claustrum surface mesh (left hemisphere); (b) Displacement is evaluated on each of the perpendicular surfaces per hemisphere, which we denote medial (M) and lateral (L). In each respective hemisphere, M is always adjacent to the putamen while L is adjacent to the insular cortex; negative displacement values on M signify inward deformation, away from the putamen, while positive values on M indicate outward deformation toward the putamen; negative values on L indicate inward deformation of the surface away from the insular cortex, while positive values indicate outward deformation toward the insular cortex; axes depicted as medial (m), lateral (l), dorsal (d), and anterior (a) show the relative direction of surface change from template vertices (*T_v_*_)_ to subject vertices (*S_v_*). (c) Thickness, computed as the Euclidean distance *d* between vertices *v_i_* and *v_j_*, related by anti-parallel, perpendicular uniform normals *n_i_* and *n_j_* connected by nearest-neighbor vertices (*k = 80*).

where the signed value *D_v_* equals the amount of variation between corresponding vertex positions of each subject’s surface (*S_v_*) and the template surface (*T_v_*), normalized by the angle-weighted outward unit normal *n̅_v_*.

Because the claustrum is a thin, closed sheet with multi-directional curvature rather than an open cortical ribbon, we treated its surface as a deformable thin shell (Terzopoulos and Fleischer 1988) to study its thickness at multiple points, rather than as two independently reconstructed surfaces as in standard cortical thickness estimation. Vertex-wise thickness was then computed as the distance between matched vertices on the shell’s two opposing faces, using the same nearest-distance approach developed for cortical thickness (Fischl and Dale 2000) and shown in Fig. 2c. Global thickness was computed from segmentation-derived claustrum volumes and mesh surface areas (SA; 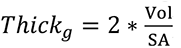).

### 2.7 Statistical Analyses

#### Global statistics

We applied a random-intercept linear mixed model (LMM) with *statsmodels.MixedLM* to investigate within-subject variances and their effects on global claustrum volume and thickness. To ensure findings were not driven by multi-site acquisition differences, claustrum volume and global thickness metrics were adjusted for multiple scanner types using the R library *longCombat* (Beer et al. 2020), a ComBat harmonization framework fine-tuned for longitudinal data. Models were evaluated both before and after scanner harmonization.

#### Marginal model

To assess longitudinal vertex-wise structural changes, we used a Generalized Estimating Equation (GEE) framework (Zeger et al. 1988). For each subject *i* with *n_i_* sessions, we specified the marginal linear model:

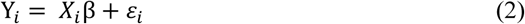

with *Y_i_* the vector of vertex measurements across sessions; *Χ_i_* the *n_i_* × *p* design matrix (including nuisance covariates); and β the *p* × 1 vector of fixed-effect parameters. This provided a computationally efficient alternative to traditional random-intercept LMMs. LMMs rely on iterative maximum likelihood estimation to model subject-specific random effects, which becomes computationally prohibitive in vertex-wise estimation and prone to failures in mass univariate settings. We therefore estimated population-averaged fixed effects with pooled ordinary least squares (OLS) across all sessions. We then applied cluster-robust (sandwich) standard errors (SE), clustering by subject to account for unequal-length longitudinal sampling (Guillaume et al. 2014). The resulting GEE framework yielded consistent parameter estimates and SE that are robust to the misspecification of the within-subject covariance structure.

#### Permutation-based cluster-wise inference

To control the Family Wise Error Rate (FWE) across the surface meshes without assuming parametric spatial smoothness, we used a non-parametric cluster-mass permutation framework. We formed t-statistics for each contrast from the OLS point estimate and subject-clustered SE, and we set a cluster-forming threshold (CFT) to *t* > 2. Vertices exceeding this CFT and connected by adjacent mesh edges were grouped into clusters; cluster-mass was the sum of *t* within each cluster (Nichols & Holmes 2002). For inference, we applied the Freedman-Lane partial permutation procedure (Freedman & Lane 1983), which isolates the primary contrast while adjusting for nuisance covariates.

Standard permutation assumes independent observations, which is violated in longitudinal data. We therefore enforced exchangeability at the subject level: we fit a reduced model omitting the term of interest, extracted the residuals, and applied a random subject-level sign flip (all sessions from a given subject received the same random ±1 flip), preserving subject correlation under the null hypothesis (Winkler et al. 2014). We then added the permuted residuals back to the reduced model’s fitted values, refit the full model, and extracted the maximum cluster-mass. Finally, we computed cluster-level p-values as the proportion of permutations whose maximum cluster-mass equaled or exceeded each observed cluster’s mass, controlling FWE across the whole surface (Nichols & Holmes 2002). This combination of marginal modeling and permutation therefore remains tractable for high-dimensional, vertex-wise, repeated measures.

#### Vertex-wise contrasts and cognitive associations

Within the full cohort, signed surface displacement and thickness were tested under the GEE framework described above, with sex and scanner model included as nuisance covariates. Three contrasts were tested independently: the group main effect at the pooled mean age (16.8), the age main effect representing the developmental slope in HC, and the *group x age* interaction.

We complemented full-cohort interaction tests with parallel within-group analyses in HC and 22q11DS. Within each group, we fit models that paralleled the full-cohort specification but replaced the group contrast with clinical/cognitive measures, testing their association with mean displacement (averaged across all vertices) using *statsmodels.MixedLM*; all continuous predictors were centered at their means. Planned secondary analyses targeted Digit Span measures (DST, DSB, DSF) as indices for working memory; FSIQ/VIQ, SST, and diagnoses of psychotic symptoms, ADHD, and lifetime anxiety disorder were treated as exploratory (Suppl. Mats. Section 4).

#### Generalized Additive Mixed Models

Because neurodevelopmental trajectories are inherently non-linear, we characterized the trajectory shape of individual clusters post-hoc using Generalized Additive Mixed Models (GAMMs). Model comparison (ΔAIC > 10, interaction *p* < 0.001) supported GAMM fit superiorly over standard LMMs for visualization.

#### Data visualization

Per-vertex inferential maps and cluster masks were rendered in 3D Slicer software (v5.6), with symmetric t-statistic scalar bounds matched to each contrast’s observed range and significant clusters shown as binary overlays across four orthogonal views (left/right, medial/lateral) to capture the asymmetric geometry. T-statistic maps were created in GIMP (v3.2.4); methods figures were created in draw.io (https://app.diagrams.net/); trajectory plots were created in R (ggplot2).

## 3 Results

### 3.1 Claustrum Volume Across Healthy Controls and 22q11DS Carriers

We initially compared claustrum volumes of healthy and clinical populations across age, group, and group x age contrasts using LMM analysis for each hemisphere (Suppl. Mats. Table 1). We observed significant decline in claustrum volume between the ages of 5–35 bilaterally (LH: β_age_ = -7.38 mm^3^/yr, *p* = 6.87e^-05^; RH: β_age_ = -6.2 mm^3^/yr, *p* = 0.0010). We observed a significant group x age interaction in the left hemisphere (β_grpXage_ = +6.51 mm, *p* = 0.007) with trend-level coefficients in the right (β_grpXage_ = +4.58 mm, *p* = 0.064). Our data showed males having significantly larger bilateral claustra (LH: β_sex_ = +199.8 mm, *p* = 5.22e^-23^; RH: β_sex_ = +205.5 mm, *p* = 4.72e^-25^), regardless of group assignment or TIV covariate inclusion. We did not observe significant group differences at the mean age after scanner harmonization (Suppl. Mats. Table 2).

### 3.2 Claustrum Thickness Across Healthy Controls and 22q11DS Carriers

We then analyzed longitudinal changes in global and vertex-wise thickness across all contrasts to understand both regional and global claustrum thickness differences.

We observed significant group differences in global thickness in the left hemisphere at the mean sample age (β_grp_ = −0.03 (22q > HC), *t* = -2.41, *p* = 0.016, α = 0.05), along with modest healthy age decline in HC (β_age_ = −0.002, *t* = -2.11, *p* = 0.034, α =0.05), but not their interaction (*p*_grpXage_ = 0.95) (Suppl. Mats. Table 3). Significant group x age effects on global thickness in the right hemisphere were observed (β_grpXage_ = −0.004, *t* = -2.68, *p* = 0.007, α = 0.05), with 22q11DS showing moderately reduced thickening across development compared to HC.

From the vertex-wise thickness analysis, we observed age-related ventral thickening and dorsal thinning in HC (ventral: c3, β = −0.015 (22q > HC), *p_FWE_* = 2.64e^-05^; c25, β = −0.0149*, p_FWE_* = 4.14e^-05^; dorsal: c20, β = +0.0098, *p_FWE_* = 0.0018). 22q11DS left claustra were thinner at the ventral pole (c3, β = −0.0055, *p_FWE_* = 0.0038) and thicker in the right dorsal-anterior claustrum (c173, β = +0.0046, *p_FWE_* = 0.002) at the mean age. Group x age interaction effects were also observed: decreased thickening in the left ventral pole of 22q11DS (c11, β = −0.0195, *p_FWE_* = 0.013) compared to HC, and preserved thickness in bilateral dorsal-anterior regions (LH: c80, β = +0.0146, *p_FWE_* = 0.024; RH: c129, β = +0.0138, *p_FWE_*= 0.003) (Suppl. Mats. Fig. 1).

### 3.3 Developmental Surface Deformation in Healthy Controls and 22q11DS Carriers

To characterize developmental change in the claustrum, we assessed surface variation by observing clusters of coordinated inward or outward deformation in each subject’s bilateral claustrum surfaces. Table 2 shows the 13 significant clusters from all contrasts. Fig. 3 shows their relative positioning and overlap along a 2D axis.

**Fig. 3.**
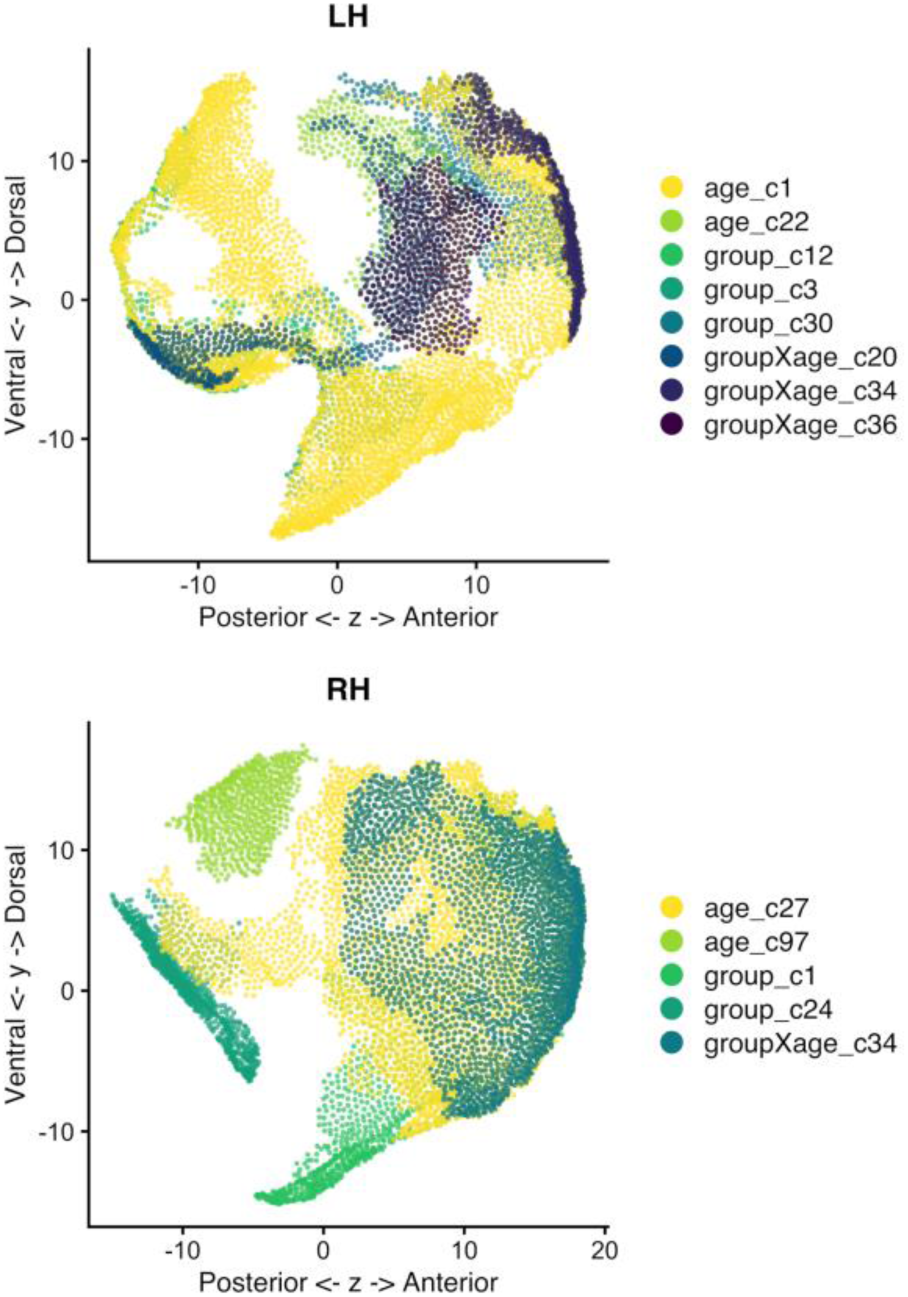
Cluster positions along the z and y axes. Individual vertices are shown with an opacity of 0.8 to show their overlap with all contrasts.

**Table 1.**
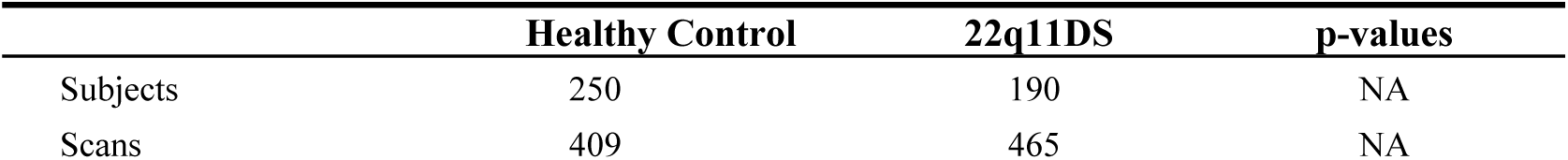
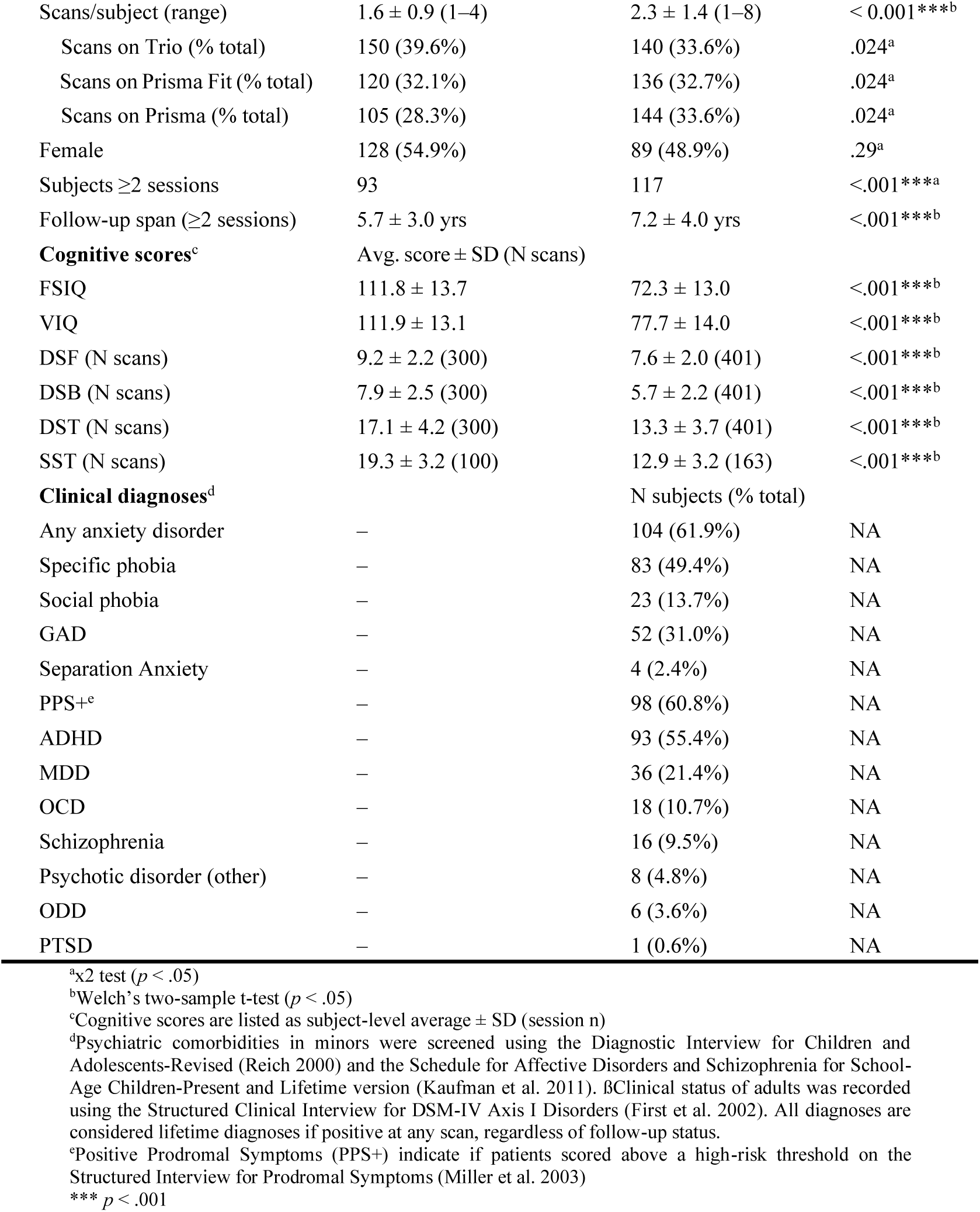
Participant demographics and cognitive scores.

**Table 2.** Displacement clusters surviving FWE-correction.

| Hemi | ID | Vertices (n) | $p_{\text{FWE}}$ | Peak $\vert t\vert $ ($\mu$) |
| --- | --- | --- | --- | --- |
| *Age* |  |  |  |  |
| L | c1 | 4,916 | **0.0002\*\*\*** | 7.90 (-0.20) |
| L | c22 | 627 | 0.0152* | 4.33 (-2.98) |
| R | c27 | 4,467 | **0.0002\*\*\*** | 9.65 (-0.40) |
| R | c97 | 634 | 0.0256* | 7.17 (-0.03) |
| *Group* |  |  |  |  |
| L | c3 | 408 | 0.0444* | 5.09 (-3.13) |
| L | c12 | 752 | 0.0046** | 7.03 (-2.91) |
| L | c30 | 736 | 0.0072** | 5.42 (+1.70) |
| R | c1 | 2,235 | **0.0002\*\*\*** | 5.15 (+0.99) |
| R | c24 | 947 | 0.0104* | 5.14 (+0.12) |
| *Group x Age* |  |  |  |  |
| L | c20 | 683 | 0.0212* | 4.73 (-2.49) |
| L | c34 | 633 | 0.0228* | 4.55 (+0.66) |
| L | c36 | 449 | 0.0488* | 4.05 (+2.82) |
| R | c34 | 2,412 | **0.0002\*\*\*** | 5.40 (+1.19) |

\*\*\* $p_{\text{FWE}} < 0.001$ ; \*\* $p_{\text{FWE}} < 0.01$ ; \* $p_{\text{FWE}} < 0.05$
Full coefficients reported in Suppl. Mats. Tables 4–6

#### Age-related deformation in healthy controls

In healthy controls, inward deformation occurred in the dorsal-anterior territory of the left medial surface, and outward deformation occurred in the same zone on the lateral surface, toward the insular cortex. Along the ventral-anterior axis, observed deformation moved perpendicularly in the opposite direction, toward the putamen, with the strongest effect at the ventral pole (c1, β = −0.0018, *t* = -3.52, *p*_FWE_ = 0.0004; Fig. 3a). In the anterior half of the right claustrum, along the dorsal-ventral axis, we observed coordinated deformation toward the insular cortex (inward medially/outward laterally; c34, β = −0.010, *t* = -7.45, *p*_FWE_ = 9.37e^-14^; Fig. 3b).

#### Cluster-mean developmental trajectories

Within each FWE-corrected group x age cluster, we assessed the average amount of surface variation per cluster using GAMMs. This allowed a focused perspective on the differences in linearity between cluster trajectories. From this analysis, we observed a progressively divergent slope across development, reflected by a characteristic plateau around age 20 (Fig. 4). This was quantified by Cohen’s d, which grew from near zero or negative in childhood (d = - 0.27 to −0.50 at age 5) to large positive values by early adulthood (d = +0.76 to +1.21 at age 35) (Suppl. Mats. Table 7).

**Fig. 4.**
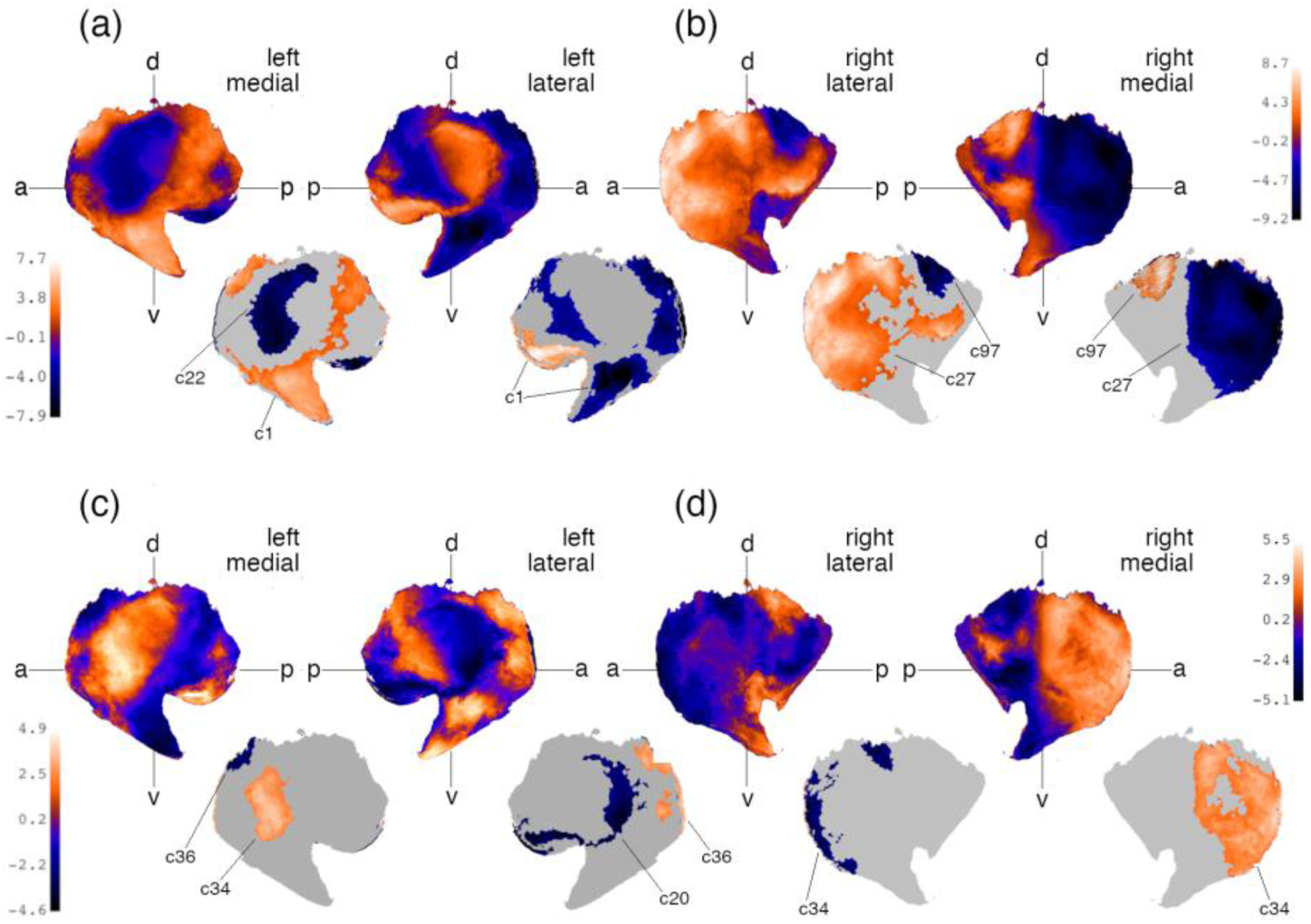
Maps of surface displacement from pooled group analysis. The age contrast is shown in (a, b). The group x age contrast (c, d) highlights regions of diagnostic group difference throughout development (22q11DS > HC); significant clusters were localized to the left dorsal-anterior claustrum (c18, *t* = -4.58, *p*_FWE_ = 0.0084; c35, *t* = +3.75, *p*_FWE_ = 0.0088; c38, *t* = +3.66, *p*_FWE_ = 0.038), while a single cluster (c34, *t* = +4.22, *p*_FWE_ = 2.42e^-05^) occupied the right anterior surface. Claustra with t-statistics are shown with labeled axes (ventral (v)–dorsal (d), anterior (a)– posterior (p)); cluster labels mark significant clusters (*p*_FWE_ < 0.05); colorbars give the t-statistic range for each hemisphere and contrast: negative values indicate inward and positive values indicate outward deformation.

**Fig. 5.**
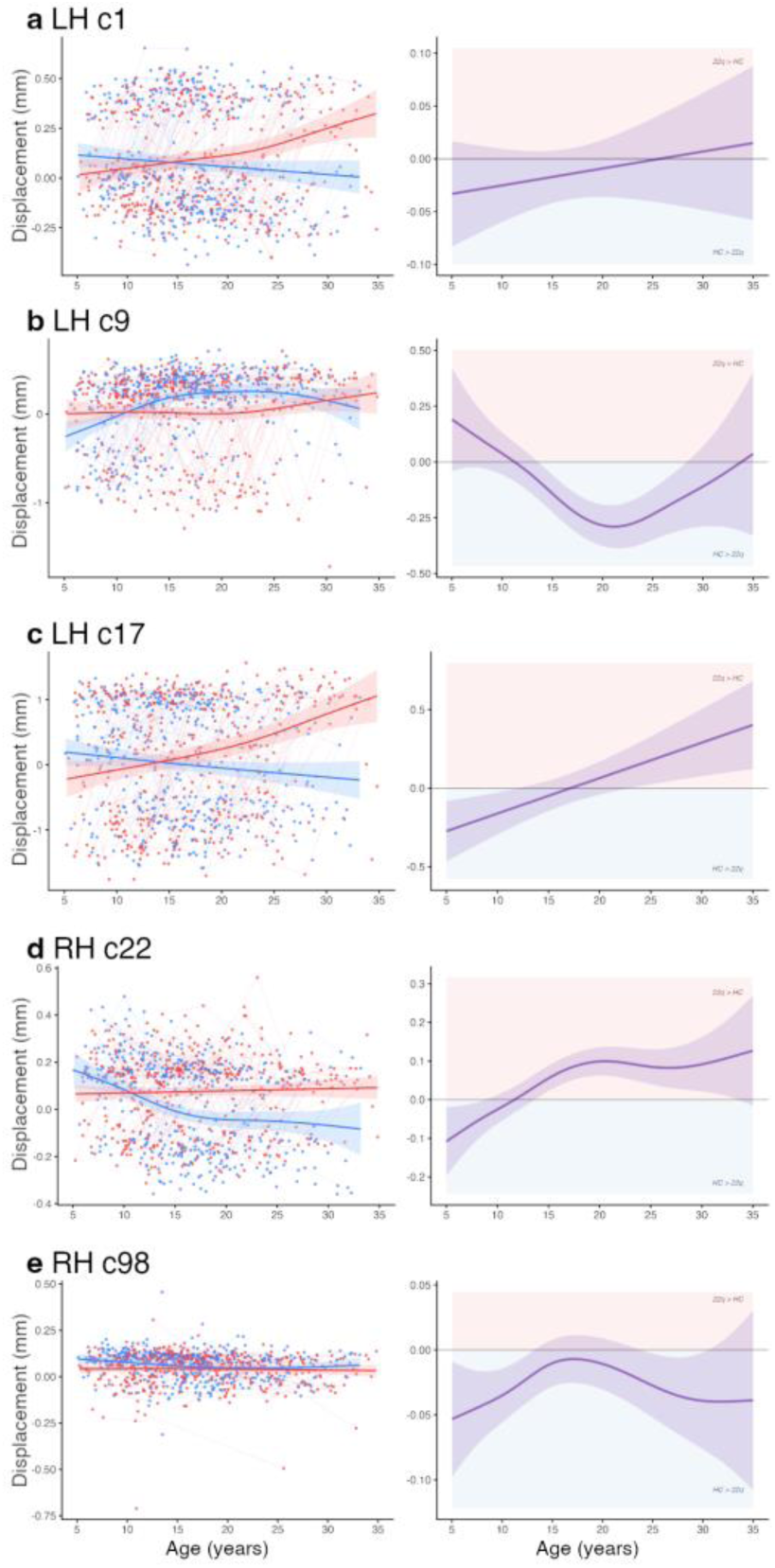
Cluster-mean trajectories of significant clusters from the group x age contrast. The left column plots displacement by age, showing nonlinear trajectories from GAMM analysis; the difference smooth line (right), shaded with a 95% CI band, shows the direction (above zero means 22q11DS carriers show higher displacement values) and the ages at which groups differ within each cluster.

One cluster in the left hemisphere (c18) showed a mean-displacement trajectory in healthy controls that rose non-linearly throughout childhood (EDF = 3.12, *p* < 0.0001, R^2^ = 0.13) and plateaued around age 15, while 22q11DS slopes showed a linear trend (EDF = 1.00, *p* = 0.135, R^2^ = 0.003), yielding a Cohen’s d that crossed from positive in childhood (+0.44 at age 5) to negative by adulthood (-1.11 at age 35) (Suppl. Mats. Table 7).

### 3.4 Cognitive Phenotyping: Working Memory and Surface-Based Measures

We next examined whether verbal and spatial working memory performance was associated with measures of claustrum surface morphology, modeling each diagnostic group separately.

#### Claustrum morphology associates with working memory in healthy controls

In HC, Digit Span scores tracked developmental slopes bilaterally (Fig. 6). Significant inward deformation of the medial surface, spanning the ventral-to-dorsal-posterior region, was associated with DSB in both the left (c3, μ(*t*) = -2.49, *p* = 0.0004; Fig. 6a) and right hemispheres (c10, μ(*t*) = -1.44, *p* = 0.018; Fig. 6b). Cluster c10 was one of the two clusters only observed in the DSB x age contrast; the other was an outwardly deforming cluster in the left lateral dorsal-posterior zone (c36, *t* = +2.89, *p* = 0.044).

**Fig. 6.**
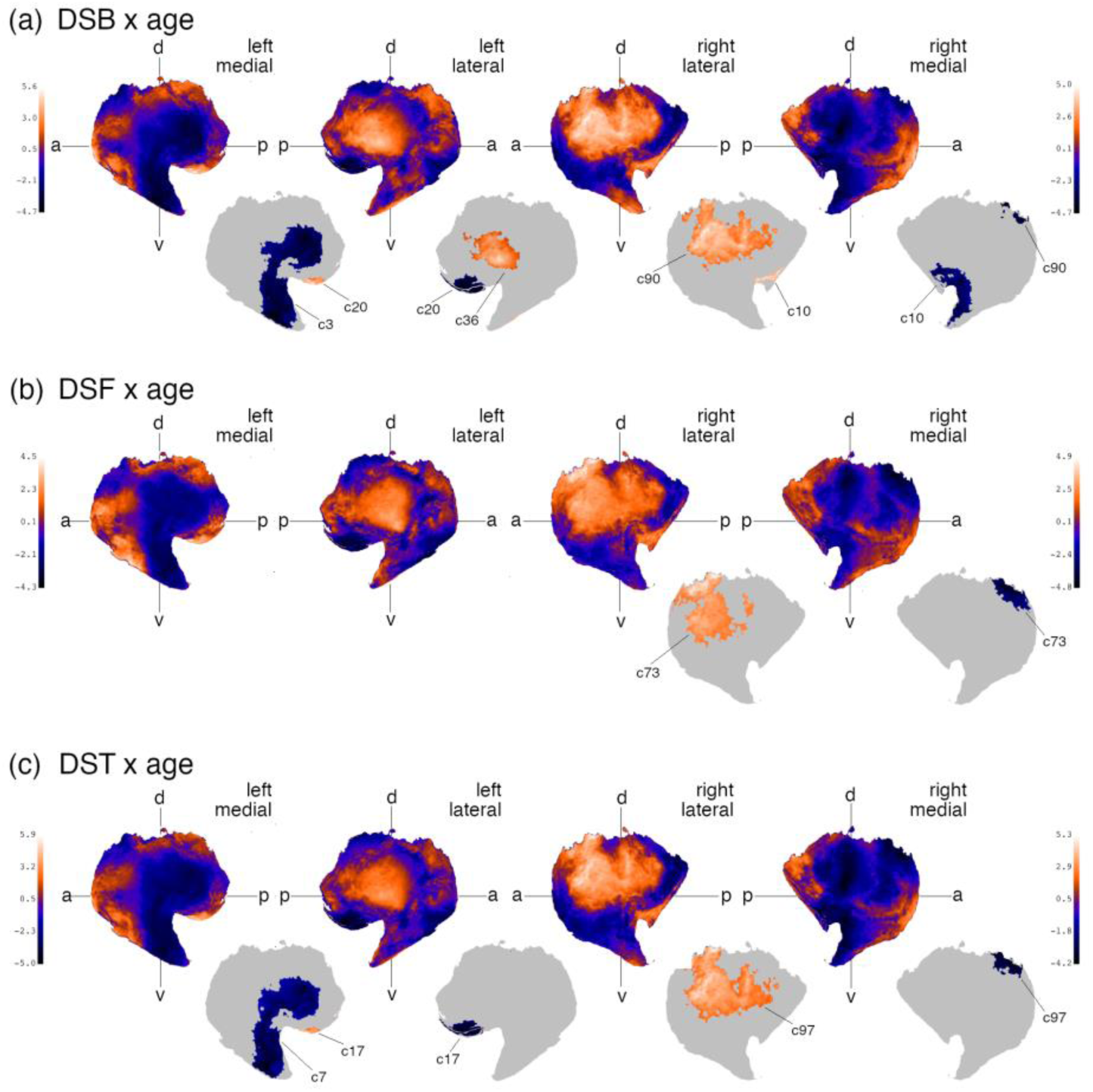
Vertex-wise t-statistic maps for the within-HC Digit Span x age interactions. shown on medial and lateral surfaces for each hemisphere. The t-statistic patterns are shown with labeled axes: dorsal (d); ventral (v); anterior (a); posterior (p); significant clusters are individually labeled for each contrast.

In one DSB x age cluster, we observed outward deformation of the right lateral surface in the dorsal-anterior region (c90, μ(*t*) = +2.49, *p* = 0.0038; Fig. 6b). In this same region, both DST x age (c97, μ(*t*) = +2.08, *p* = 0.0038) and DSF x age (c73, μ(*t*) = +1.22, *p* = 0.018) interactions associated with surface deformation significantly.

#### Claustrum morphology associates uniquely with verbal and visuo-spatial working memory

Claustrum surface deformation patterns across ages 5–35 and their relation to verbal (DST) and visuo-spatial (SST) working memory tests are shown in Fig. 7. We observed differing patterns across test domains by the spatial localization and magnitude of their t-statistics. The DST x age contrast showed inward deformation of the left medial surface that was distributed posteriorly along the dorsoventral axis, while the SST effect was distributed anteriorly and deformed outward in relation to age (Fig. 7a, b). The right lateral surface deformation effect was concentrated along the anterior edge in the SST contrast, whereas the DST contrast effect was concentrated dorsally. No significant clusters were observed in the SST model after FWE-correction.

**Fig. 7.**
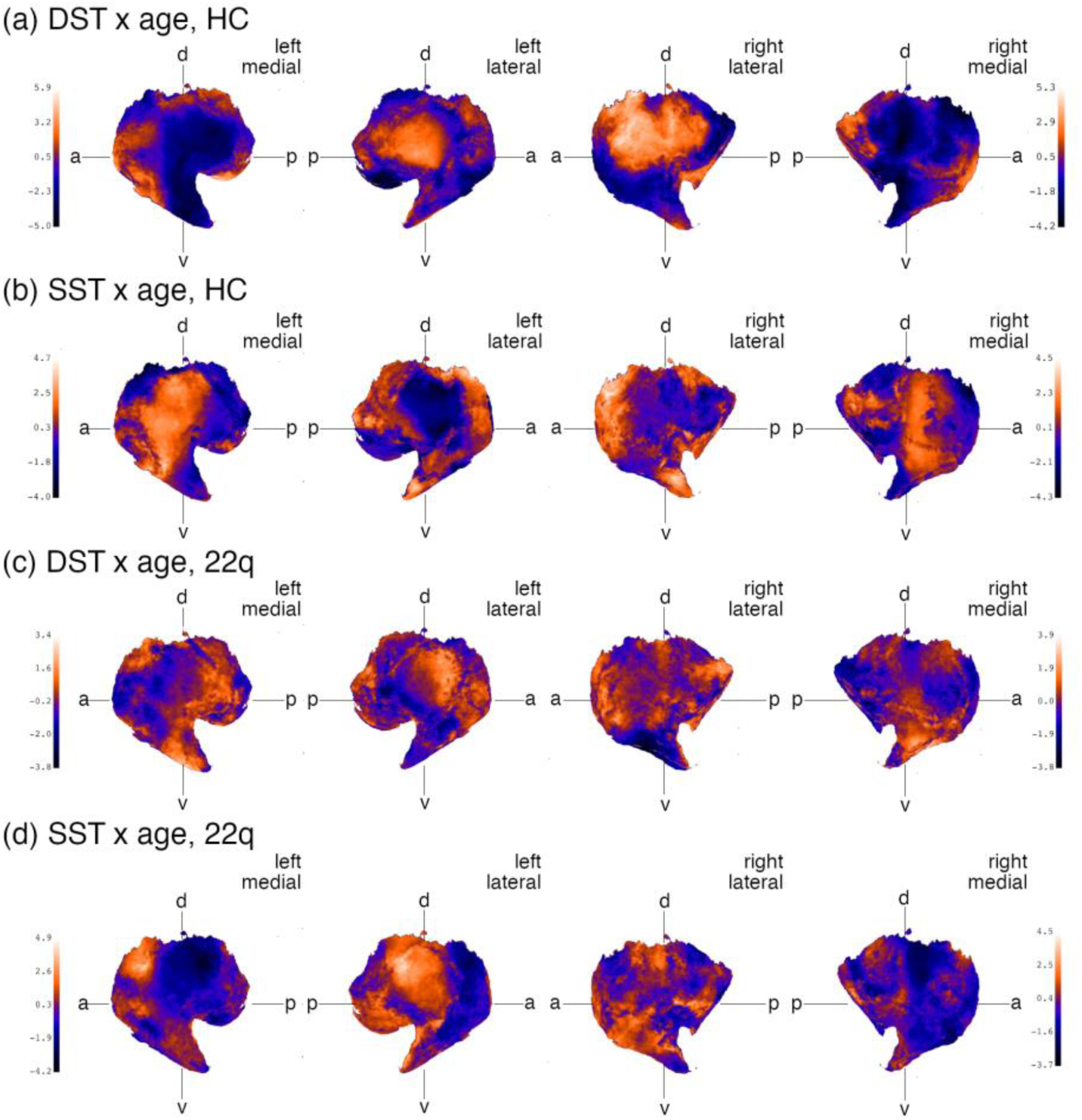
Comparison of Digit and Spatial Span displacement t-statistics between groups. The within-HC model results for DST x age (a) and SST x age (b) results are shown per hemisphere; within-22q11DS results are shown in (c) and (d).

**Fig. 8.**
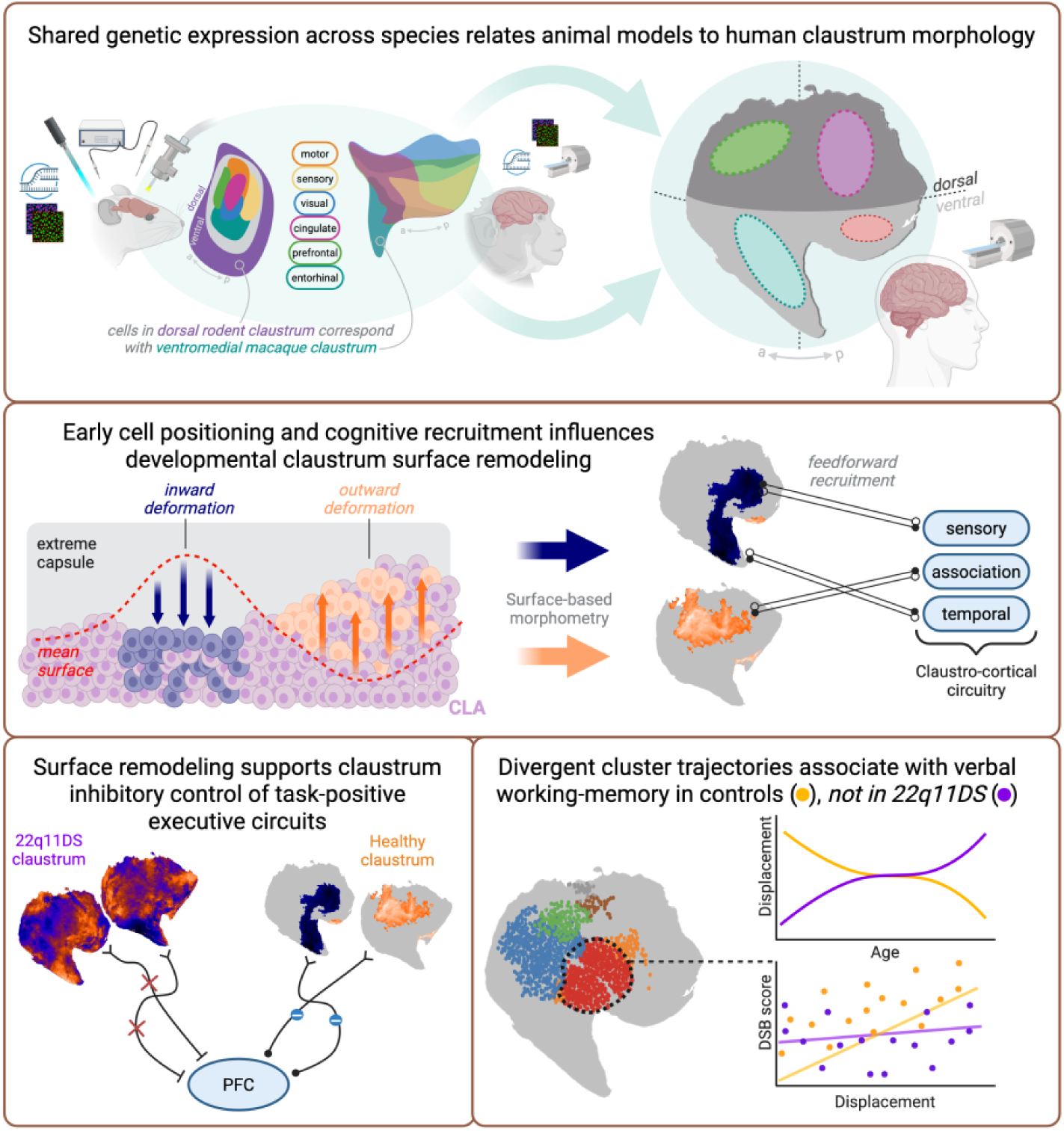
Diagram showing the hypotheses derived from surface morphometry. Shared cellular makeup in the claustrum complex allows the comparison of projection topography between animal and human models (top). Alterations in cell positioning and cognitive recruitment in early developmental stages are hypothesized to lead to structural differences in healthy vs 22q11DS carrier claustra (middle). The organization of deforming surface vertices is hypothesized to relate to 22q11DS-specific working memory deficits (bottom). Created in https://BioRender.com.

Maps of t-statistic effects from the within-22q11DS model show a similar distinction between working memory task domains and surface deformation. The strongest difference can be observed in the left hemisphere (Fig. 7c, d), where outward deformation is seen in the ventral and ventral-posterior zones of the DST contrast, but is opposed by inward deformation on the left lateral dorsal claustrum in the SST contrast.

#### Claustrum morphology and working memory relationship is blunted or absent in 22q11DS

The within-22q11DS model differed from HC in both magnitude and spatial distribution when compared using the same covariates and cognitive predictors (Fig. 7). No significant clusters were observed in the within-22q11DS model for any Digit or Spatial Span predictor.

To evaluate whether the absence of a correlation in 22q11DS represented a true null effect rather than insufficient statistical power, we first conducted vertex-wise Two One-Sided Tests (TOST; Δ = ± 0.10) (Lakens 2017). When comparing the DSB x age coefficients between the individual within-group models across the entire claustrum surface (all 10,000 vertices), 100% of vertices were statistically equivalent to zero (*p*_TOST_ < 0.05), with an overall mean effect size of β = 0.0008 (90% CI [-0.0055, +0.0071]). When restricting the analysis to the exact vertices in the HC-only significant clusters (N = 1,710 vertices), the same vertices in 22q11DS remained equivalent to zero (*p*_TOST_ < 10e^-60^ across all vertices). The absent association was not further explained by measurement range, participant age, or antipsychotic exposure (Suppl. Mats Section 5).

## 4 Discussion

We investigated claustrum development longitudinally using both volume and surface-based metrics in healthy controls and 22q11DS carriers. Our findings reveal a spatially organized developmental pattern of claustrum morphology that global volume alone does not capture. Importantly, regional claustrum morphology was associated with verbal and spatial working memory performance in healthy controls, whereas this structure–behavior relationship was absent in 22q11DS. Together, these findings point to a disruption of the typical developmental organization linking claustrum structure to cognitive function. In this section we discuss our findings in light of translational and human literature.

### 4.1 Developmental Remodeling of the Claustrum in Typical Development and 22q11DS

Our findings suggest that claustrum development involves regionally organized morphological remodeling that global measures only partly capture. Although claustrum volume declined bilaterally across the 5–35-year age range, the more informative developmental differences emerged from the regional thickness and surface analyses. In healthy controls, thickness changed in a spatially heterogeneous manner across development, whereas these regional trajectories were attenuated in 22q11DS. This pattern is consistent with lifespan MRI studies showing that subcortical maturation is region- and hemisphere-specific rather than spatially uniform (Raznahan et al. 2014; Wierenga et al. 2014; Miletić et al. 2022). Altered maturational remodeling may therefore contribute to the thickness abnormalities observed in 22q11DS. Synaptic pruning provides one possible biological framework, given previous evidence of altered pruning and cortical thickness trajectories in 22q11DS (Campbell et al. 2009; Schaer et al. 2009; Mancini et al. 2020; Bagautdinova et al. 2021; Alvino et al. 2026; Pascucci et al. 2026), although T1-weighted morphometry cannot identify the cellular processes underlying macroscopic change.

Surface displacement revealed a related but qualitatively different abnormality. Whereas thickness differences largely reflected attenuation of the typical developmental trajectory, displacement trajectories diverged in direction between groups in several regions. The 22q11DS phenotype is therefore not simply a weaker version of normative maturation. These differences were present from the youngest age examined, suggesting that atypical claustral organization is established before, or early within, the developmental period sampled here and then evolves along a different trajectory. The present data cannot determine when such alterations arise, or which biological process produces them.

One possible framework involves early differences in neuronal positioning and claustral organization. Claustrum formation depends on tightly coordinated neuronal migration and molecular specification, and perturbing these processes alters claustral shape and connectivity in animal models: Nr4a2 deficiency displaces claustral neurons toward the insular cortex and modifies claustral morphology (Mantas et al. 2024; Yan et al. 2026), and disrupted neocortical development produces cytoarchitectural deficits such as altered cell positioning (Gangatharan et al. 2018; Oshima et al. 2023), a mechanism well-documented in psychiatric disorders (Jakob and Beckmann 1986; Beckmann and Jakob 1991; Casanova et al. 2008; Kitazawa et al. 2014). These findings offer a plausible model by which early structural organization could shape later surface trajectories. However, no comparable mechanism has been demonstrated in humans or in 22q11DS; the present morphometric findings generate a hypothesis about neuronal migration, not an explanation established by them.

### 4.2 Verbal Working Memory Performance and Claustrum Surface Morphology in Healthy Controls and 22q11DS

In healthy controls, higher Digit Span task scores were associated with greater surface displacement in ventral-posterior and dorsal-anterior clusters. DSB correlated uniquely with clusters in the ventral and dorsal-posterior zones, whereas DSF, an index of simple storage without manipulation, related only to surface variation in the dorsal-anterior claustrum. Because DSB, but not DSF, selectively engages the dlPFC (Owen et al. 1996, 2002) and dorsal ACC (Duncan and Owen 2000), deformation within these zones may form part of the structural substrate through which the claustrum coordinates executive and sensory networks during verbal working memory maintenance. The same areas were not significantly deformed in SST x age analyses, suggesting that deformation in the ventral-posterior zone associates more with verbal than visuo-spatial working memory. The within-group models showed SST x age deformation distributed anteriorly, while DST x age effects were organized dorsally or posteriorly.

Building on this anatomical correspondence, the ventral-posterior DSB-related cluster may be particularly relevant to interactions between the claustrum, DMN, and executive networks during working memory maintenance. In healthy controls, higher DSB scores were associated with greater inward deformation across age in this region. During working memory maintenance, DMN activity is typically reduced relative to task-positive networks (Piccoli et al. 2015); consistent with this, DMN deactivation during maintenance aligns with the direction of ventral deformation in HC. In 22q11DS, the DSB–morphology association was absent, and the developmental deformation trajectory differed in direction, suggesting that this typical structure– behavior relationship is disrupted.

The relevance of this association to working memory is further supported by its relative specificity across the cognitive measures examined. The DSB-displacement relationship persisted after adjustment for FSIQ and VIQ. This pattern converges with functional evidence that claustrum engagement accompanies DMN deactivation during working memory tasks, rather than during internally directed cognition more broadly (Huang et al. 2025; Stewart et al. 2026). Although attention contributes to DSB performance (Hale et al. 2002), ADHD diagnosis did not mediate the DSB-displacement association in our data, indicating that the relationship is not a generic correlate of executive function or attention.

### 4.3 Working Memory and Claustrum Surface Morphology Are Not Associated in 22q11DS

In 22q11DS, claustrum morphology was not associated with Digit or Spatial Span performance. One interpretation is that the association observed in controls may itself emerge through maturation. Experience-dependent plasticity shapes claustro-frontal circuits (Chaudhury et al. 2016; Hansel and Yuste 2024), and claustro-cortical projections to the ACC engage PV interneurons that switch from excitation to inhibition within a precise developmental window (Shaker et al. 2026). PV-mediated inhibition could thus gate the maturation of these circuits, tuning claustral structure to individual differences in working memory performance. If this maturation fails or is delayed in 22q11DS, claustral morphology would remain untuned to verbal or visuo-spatial working memory ability, which may explain the null morphology-behavior association.

This proposed maturational difference may also have a spatial expression. In healthy controls, the morphology-working memory relationship was regionally differentiated, with verbal tasks relating primarily to ventral-posterior territories and visuospatial tasks to more anterior and dorsal regions. This spatial differentiation was not evident in the right claustrum of 22q11DS, where DST and SST showed more overlapping deformation patterns. If the regional organization of these associations partly reflects maturation of distinct claustro-cortical circuits, its loss in 22q11DS could represent another manifestation of atypical circuit development. However, such an interpretation remains speculative because surface morphology alone cannot determine whether these patterns reflect altered connectivity, inhibitory maturation, or another developmental mechanism. Task-based claustrum imaging in humans, complemented by causal circuit studies in animal models, will be needed to distinguish among these possibilities.

## 5 Strengths, Methodological Considerations, and Future Directions

Overall, we show that analyzing claustrum surface morphology complements simple volumetric studies, capturing changes in more complex surface metrics across development. Using this methodology, we identified regional territories associated with verbal working memory performance in typically developing individuals, but not in 22q11DS carriers. Early alterations in claustral organization and later abnormalities in circuit maturation are not mutually exclusive: atypical early organization could provide a different structural substrate for later experience-dependent and inhibitory maturation. Our results suggest that developmental remodeling of the claustrum surface may reflect maturation of network-level claustral circuits; disruption of these mechanisms could contribute to the working memory vulnerability of 22q11DS.

We acknowledge several limitations of this study. Our longitudinal template construction is affected by site, scanner, and protocol drift, which confounds sub-millimeter structures like the claustrum more than larger regions, compounded by uneven sampling density across the age range. A second limitation is distinguishing true shape change from repositioning due to the neighboring external and extreme capsules (Mathur et al. 2009), which requires an independent estimate of those boundaries, which we lack. This is especially relevant in 22q11DS where architectural anomalies in these regions are documented (Kikinis et al. 2012; Tylee et al. 2017; Villalón-Reina et al. 2020), and may themselves be age-dependent, bearing directly on our group x age results. Surface-based analysis is further limited by the claustrum’s thinness relative to structural T1w MRI resolution, where partial volume effects, surface-correspondence assumptions, and claustrum-insula border ambiguity are unavoidable consequences of segmentation. Automated pipelines also inherit systematic claustrum-insula-putamen mislabeling from their deep-learning training labels (Valabregue et al. 2024). Finally, this study lacks a direct neural measure of working memory, so we cannot establish causality between morphological changes and behavior.

These limitations point to several tractable next steps: diffusion tractography of the extreme and external capsules to delineate claustrum boundaries independently of the segmentation surface; deep-learning segmentation trained on high-resolution 7T data, which has begun to improve claustrum delineation (Li et al. 2021; Albishri et al. 2022; Coates and Zaretskaya 2024); task-based fMRI to test directly whether claustral functional connectivity tracks working memory performance and whether or not this is due to disorganized deformation patterns in 22q11DS; and translational work linking the affected surface territories identified here to genetically defined claustral cell populations in animal models, to test whether these effects relate to the activation of specific projection neuron subtypes.

In conclusion, this study shows that claustrum maturation is spatially heterogeneous and that regional surface morphology captures developmental changes obscured by global volumetry. The association between claustrum morphology and working memory performance in healthy controls, and its disruption in 22q11DS, further link regional claustrum maturation to cognitive development and highlight altered structure–behavior coupling in neurodevelopmental dysmaturation. More broadly, these findings support longitudinal surface-based shape analysis as a sensitive framework for resolving developmental changes in small, anatomically heterogeneous subcortical structures.

## Author contributions

AA: Conceptualization, Methodology, Software, Formal analysis and investigation, Writing - original draft preparation, Writing - review and editing. FD: Conceptualization, Formal analysis and investigation, Writing -original draft preparation, Writing - review and editing, Supervision. CL: Clinical insight, Writing - review and editing. DVDV: Conceptualization, Methodology, Writing - review and editing, Resources, Supervision. SE: Conceptualization, Writing - review and editing, Resources, Funding acquisition, Supervision.

## Acknowledgments.

This work was made possible by the generous support of the Fondation Privée des HUG, the Fondation Coplay-May, and Fondation Pôle Autisme, Geneva, Switzerland. Their commitment to advancing research enabled the results presented here.

## Data availability

The datasets generated during and/or analyzed during the current study are not publicly available due to confidentiality but anonymized dataset are available from the corresponding author on reasonable request.

## Ethical approval

The study was approved by the ethics committee for research (CCER) and conducted according to the Declaration of Helsinki.

## Consent

Written informed consent was obtained from either participants or their parents (for minors).

## Declaration of generative AI and AI-assisted technologies in the manuscript preparation process

During the preparation of this work, the authors used Anthropic’s Claude (Opus 4.8) to filter demographic data in the preprocessing stages and the Claude skill *academic-humanizer* (https://github.com/AIScientists-Dev/academic-humanizer) was used by the same model for tuning the academic style of original writing. After using this tool/service, the authors reviewed and edited the content as needed and take full responsibility for the content of the published article. and for the organization of ideas during manuscript preparation.

