## supplementary_materials for "Claustrum Morphometry Reveals Spatially Heterogeneous Alterations in 22q11DS"

### 1 Quality control and exclusions

#### 1.1 Scanner distortion

Cluster-mean displacement at several FWE-significant clusters showed a visually striking per-scanner intercept offset of approximately ±1.5–2 mm (Fig. 1, 2). Within-scanner age slopes at every such cluster are parallel and same-signed, and the LMM scanner factor absorbs the intercept offset before the tested coefficients are estimated. To further elucidate that this offset does not reflect a pipeline artifact or a bias in the inferential results, we ran two diagnostics: one characterizing the structure of the offset, and one showing its spatial overlap with the effect maps. Effectively, we show how the visible offset has the signature of a smooth gradient distortion field.

For each pair of scanners and each hemisphere, we computed the per-vertex mean displacement difference across all sessions on each scanner (Table 1). Spatial autocorrelation of change per scanner was quantified by Moran's I on the grand template mesh adjacency graph, with a 1,000-permutation null distribution generated by random vertex shuffling.

All six maps showed Moran's I between +0.83 and +0.91, far above any permutation null value, indicating that the offset is a smooth, spatially coherent field rather than uncorrelated per-vertex noise. This pattern is the diagnostic signature of regional gradient nonlinearity and B0 inhomogeneity (Jovicich et al. 2006; Wang et al. 2004), which produce position-dependent geometric displacements on the order of 1–2 mm at

3–4 cm of isocenter on 3T Siemens systems. The spatial structure and magnitude both match the observed change per scanner. The hypothesis that the offset reflects a pipeline bug, segmentation drift, or template instability is incompatible with this level of spatial coherence.

**Table 1.** Per-scanner distortion metrics.

| Hemi | Pair | Mean Δ (mm) | SD | Range | Moran's I | *p* |
|---|---|---|---|---|---|---|
| LH | Prisma − Prisma_fit | +0.32 | 1.05 | [−3.4, +4.2] | **+0.914** | < .001 |
| LH | Prisma − Trio | +0.38 | 1.28 | [−4.9, +4.9] | **+0.905** | < .001 |
| LH | Prisma_fit − Trio | +0.06 | 0.36 | [−1.6, +1.7] | **+0.877** | < .001 |
| RH | Prisma − Prisma_fit | −0.01 | 1.14 | [−3.7, +3.6] | **+0.829** | < .001 |
| RH | Prisma − Trio | +0.04 | 1.38 | [−3.9, +3.8] | **+0.861** | < .001 |
| RH | Prisma_fit − Trio | +0.04 | 0.38 | [−1.1, +1.3] | **+0.914** | < .001 |

a)

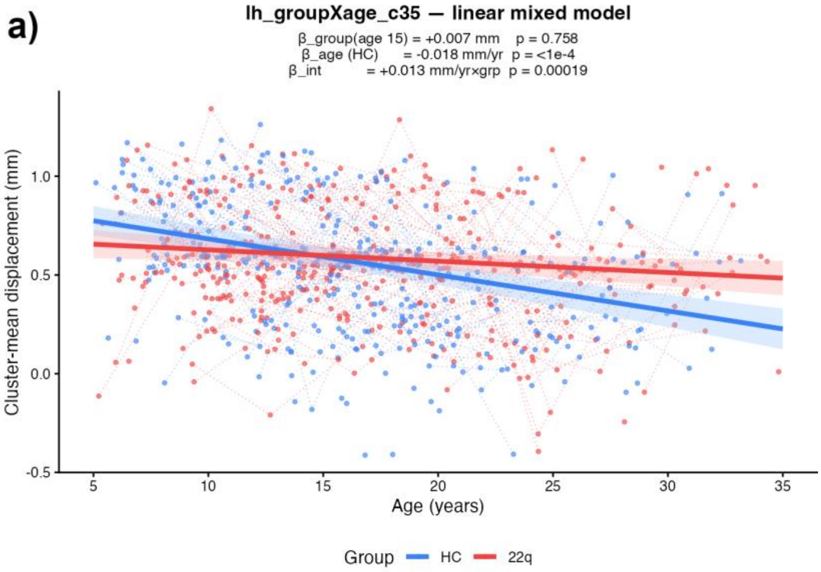
lh_groupXage_c35 — linear mixed model
β_group(age 15) = +0.007 mm p = 0.758
β_age (HC) = -0.018 mm/yr p = <1e-4
β_int = +0.013 mm/yr×grp p = 0.00019
Cluster-mean displacement (mm)
Age (years)
Group HC 22q

b)

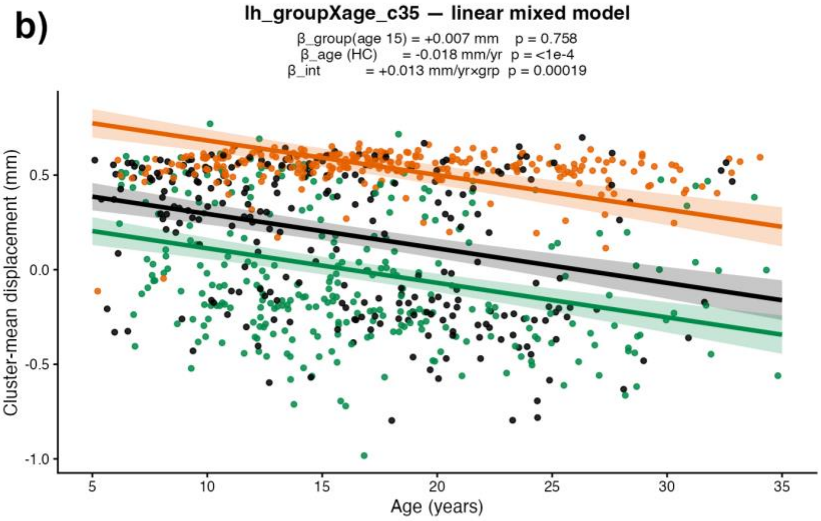
lh_groupXage_c35 — linear mixed model
β_group(age 15) = +0.007 mm p = 0.758
β_age (HC) = -0.018 mm/yr p = <1e-4
β_int = +0.013 mm/yr×grp p = 0.00019
Cluster-mean displacement (mm)
Age (years)

**Fig. 1.** Signal variability between three Siemens 3T scanners.

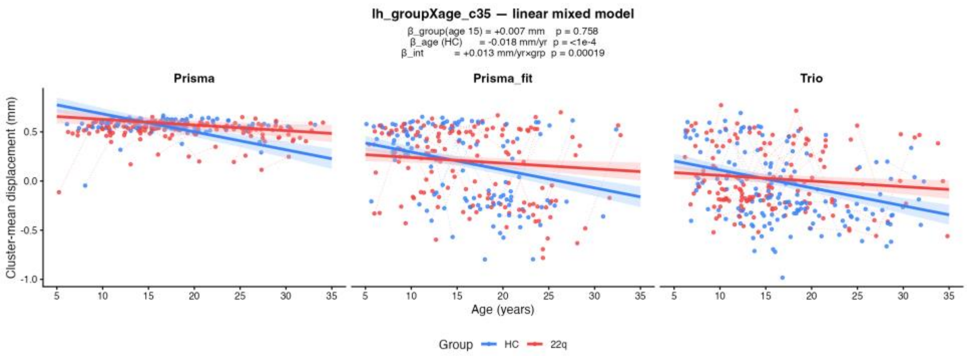


**Fig. 2.** Signal variability between three Siemens 3T scanners.

The analysis also showed spatially dissociable group-difference effects from scanner offset. For each FWE-tested main contrast (group, age, group x age) and each pairwise-offset map, we computed two measures of spatial overlap between the inferential effect and the offset across all 10,000 vertices: the Pearson correlation between $|t_{obs}|$ and change per scanner, and the Jaccard index between thresholded masks (Table 2). Group main effects in both hemispheres were uncorrelated with the scanner-offset spatial pattern.

The proportion of effect-mask vertices falling in high offset zones was at chance for both group main effects (~46% LH, ~47% RH, vs ~50% expected under spatial independence) and within or near chance for group x age (~34–77%). This dissociates the primary group inferential target from the scanner offset spatial pattern: group difference clusters live at vertices that are not preferentially located where the offset layer is largest.

**Table 2.** Spatial overlap between inference and offset.

| Hemi | Contrast | Pearson *r* | Jaccard |
|---|---|---|---|
| LH | group | −0.06 to −0.05 | 0.20 – 0.22 |
| RH | group | −0.05 to −0.02 | 0.21 – 0.22 |
| LH | age | +0.15 to +0.48 | 0.45 – 0.54 |
| RH | age | −0.16 to +0.61 | 0.33 – 0.61 |
| LH | group × age | +0.05 to +0.38 | 0.24 – 0.36 |
| RH | group × age | −0.26 to +0.39 | 0.12 – 0.27 |

### 2 Claustrum Global Volume and Thickness Across Healthy Controls and 22q11DS Carriers

#### 2.1 TIV covariation

**Table 3.** Volumetric LMM results after scanner harmonization, with and without TIV covariate inclusion.

| | No TIV covariate | | TIV covariate | |
|---|---|---|---|---|
| | **LH** | **RH** | **LH** | **RH** |
| **n_sessions** | 877 | 877 | 877 | 877 |
| **n_subjects** | 454 | 454 | 454 | 454 |
| **beta_group** | -24.644 | 24.444 | -39.372 | -3.750 |
| **se_group** | 20.470 | 20.055 | 28.571 | 28.020 |
| **p_group** | 0.229 | 0.223 | 0.168 | 0.894 |
| **beta_age** | -7.443 | -6.251 | -7.392 | -6.215 |
| **p_age** | **0.000** | **0.001** | **0.000** | **0.001** |
| **beta_groupXage** | 6.682 | 4.752 | 6.553 | 4.661 |
| **p_groupXage** | **0.006** | 0.056 | **0.007** | 0.060 |
| **beta_sex_M** | 203.121 | 208.690 | 183.651 | 177.443 |
| **p_sex_M** | **0.000** | **0.000** | **0.000** | **0.000** |
| **beta_TIV** | - | - | 40.281 | 37.006 |
| **p_TIV** | - | - | **0.000** | **0.002** |
| **sigma_u** | 183.402 | 173.478 | 182.567 | 172.681 |
| **sigma_e** | 142.530 | 152.715 | 141.386 | 151.679 |
| **icc_subject** | 0.623 | 0.563 | 0.625 | 0.564 |
| **d_age5** | -0.446 | -0.137 | -0.505 | -0.256 |
| **d_age10** | -0.302 | -0.034 | -0.363 | -0.154 |
| **d_age15** | -0.158 | 0.069 | -0.222 | -0.053 |
| **d_age20** | -0.014 | 0.172 | -0.080 | 0.049 |
| **d_age25** | 0.130 | 0.274 | 0.062 | 0.150 |
| **d_age30** | 0.274 | 0.377 | 0.204 | 0.251 |
| **d_age35** | 0.417 | 0.480 | 0.346 | 0.353 |

All beta and SE units are in $mm^3$/year

#### 2.2 Scanner harmonization

**Table 4.** Volumetric LMM results before scanner harmonization. Analysis was made for respective claustra in each hemisphere (LH/RH) and for bilateral claustra. $Age^2$, group x age, group x sex, and sex x age covariates were not significant and excluded from the final model. Cohen's d was computed at 5-year intervals in the 5–35 range. Age, sex, scanner, and TIV effects showed significance ($p < 0.05$) and were included in the final model.

| | LH | RH | Bilateral |
|---|---|---|---|
| **N_scans** | 840 | 840 | 840 |
| **N_subjects** | 434 | 434 | 434 |
| **beta_group** | 17.234 | 53.565 | 71.419 |
| **se_group** | 22.852 | 21.192 | 39.612 |
| **p_group** | 0.451 | 0.011 | 0.071 |
| **beta_age** | -5.551 | -3.395 | -8.692 |
| **p_age** | **0.002** | **0.055** | **0.008** |
| **beta_age²** | 0.067 | 0.183 | 0.257 |
| **p_age²** | 0.559 | 0.112 | 0.219 |
| **beta_groupXage** | 2.310 | 1.262 | 3.292 |
| **p_groupXage** | 0.239 | 0.511 | 0.351 |
| **beta_sex_M** | 73.899 | 46.260 | 119.883 |
| **p_sex_M** | **0.001** | **0.030** | **0.003** |
| **beta_groupXsex** | -7.938 | 27.935 | 18.829 |
| **p_groupXsex** | 0.805 | 0.348 | 0.735 |
| **beta_sexXage** | 0.938 | -1.112 | -0.344 |
| **p_sexXage** | 0.622 | 0.553 | 0.920 |
| **beta_scanner_Trio** | -28.891 | -29.590 | -58.230 |
| **p_scanner_Trio** | **0.023** | **0.021** | **0.012** |
| **beta_scanner_Prisma** | -49.323 | -67.709 | -117.647 |
| **p_scanner_Prisma** | **0.0000770** | **0.0000001** | **0.0000003** |
| **beta_TIV** | 1132.313 | 1239.859 | 2380.639 |
| **p_TIV** | **0.000000** | **0.000000** | **0.000000** |
| **sigma_u** | 137.181 | 118.436 | 227.658 |
| **sigma_e** | 115.650 | 122.578 | 218.559 |
| **icc_subject** | 0.585 | 0.483 | 0.520 |
| **d_age5** | -0.056 | 0.227 | 0.103 |
| **d_age10** | 0.009 | 0.264 | 0.155 |
| **d_age15** | 0.073 | 0.301 | 0.208 |

| | | | |
|---|---|---|---|
| **d_age20** | 0.137 | 0.338 | 0.260 |
| **d_age25** | 0.202 | 0.375 | 0.312 |
| **d_age30** | 0.266 | 0.412 | 0.364 |
| **d_age35** | 0.330 | 0.449 | 0.416 |

All beta and SE units are in $mm^3$/year

#### 2.3 Thickness

**Table 5.** Claustrum global thickness after scanner harmonization.

| Hemi | Contrast | beta | SE | t | p | CI95 | $r^2$_partial | ICC |
|---|---|---|---|---|---|---|---|---|
| Left | group | -0.03047 | 0.01264 | -2.41 | 0.0159 | [-0.05525, -0.00570] | 0.1059 | 0.49 |
| Left | age | -0.00237 | 0.00112 | -2.11 | 0.0345 | [-0.00457, -0.00017] | 0.0876 | 0.49 |
| Left | groupXage | $9.00e^{-05}$ | 0.00143 | 0.06 | 0.9521 | [-0.00273, +0.00290] | 0.0902 | 0.49 |
| Right | group | 0.00118 | 0.01446 | 0.08 | 0.935 | [-0.02717, +0.02953] | 0.0721 | 0.59 |
| Right | age | 0.00247 | 0.00121 | 2.04 | 0.0411 | [+0.00010, +0.00483] | 0.0747 | 0.59 |
| Right | groupXage | -0.00415 | 0.00155 | -2.68 | 0.0074 | [-0.00719, -0.00111] | 0.077 | 0.59 |

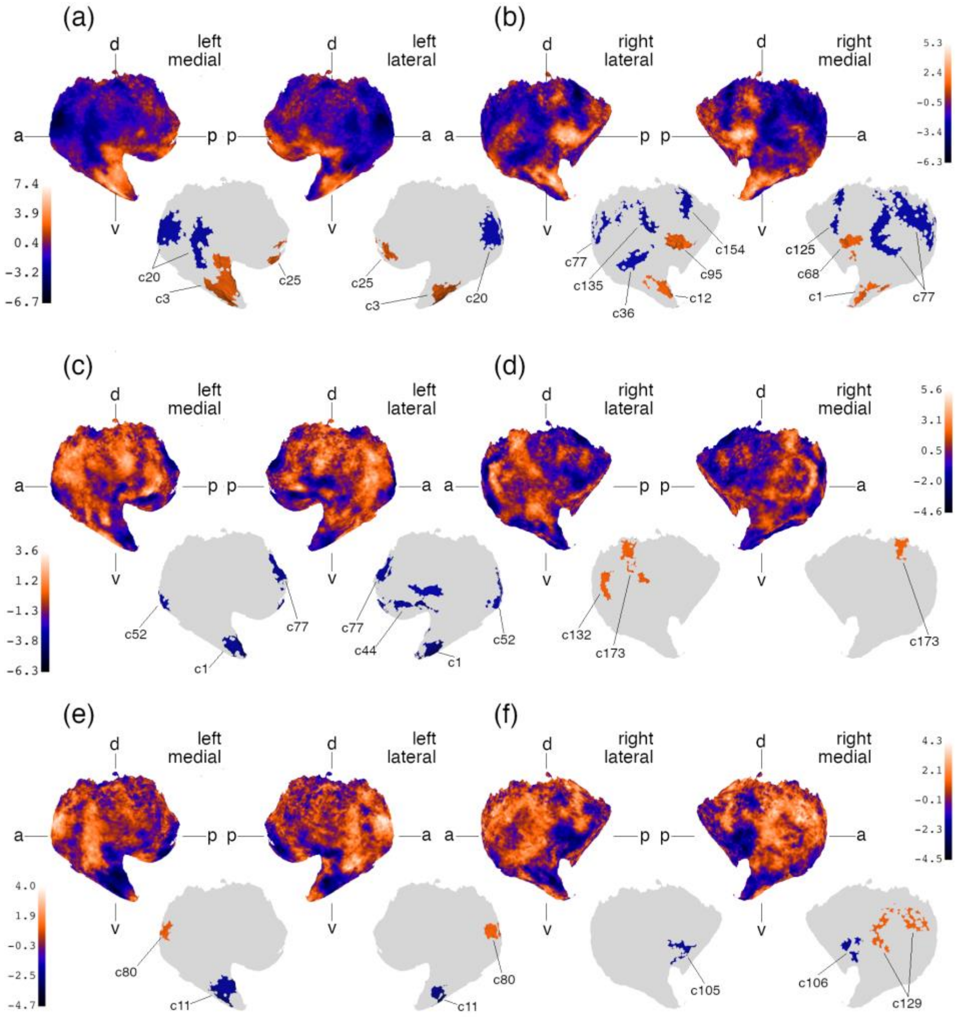


**Fig. 3.** Vertex-wise t-statistic maps for claustrum thickness across the age (a, b), group (c, d), and group-by-age (e, f) contrasts, shown on medial and lateral surfaces for each hemisphere (LH: a, c, e; RH: b, d, f). Colorbars give the t-statistic range for each panel. Lower insets show clusters surviving correction for each contrast, colored by t-statistic sign. d, dorsal; v, ventral; a, anterior; p, posterior.

### 3 Developmental Surface Deformation in Healthy Controls and 22q11DS Carriers

**Table 6.** FWE-corrected displacement clusters, age contrast.

| ID | c1 | c22 | c27 | c97 |
|---|---|---|---|---|

| **Hemi** | L | L | R | R |
|---|---|---|---|---|
| **Vertices (n)** | 4,916 | 627 | 4,467 | 634 |
| **$p_{FWE}$** | 0.0002* | 0.0152 | 0.0002* | 0.0256 |
| **Peak \|t\| (μ)** | 7.90 (-0.20) | 4.33 (-2.98) | 9.65 (-0.40) | 7.17 (-0.03) |
| **age (β, p)** | -0.002, .1548 | -0.022, .0003 | -0.01, 1.5e$^{-11}$ | -0.002, .0179 |
| **group (β, p)** | -0.026, .0177 | -0.024, .635 | +0.043, 1.56e$^{-05}$ | -0.015, .0024 |
| **group x age (β, p)** | -0.001, .711 | +0.023, .0023 | +0.006, .0001 | +0.001, .553 |
| **sex (β, p)** | +0.034, .0018 | -0.028, .562 | +0.010, .325 | +0.010, .064 |

**Table 7.** FWE-corrected displacement clusters, group contrast.

| **Hemi** | L | L | L | R | R |
|---|---|---|---|---|---|
| **ID** | c3 | c12 | c30 | c1 | c24 |
| **Vertices (n)** | 408 | 752 | 736 | 2,235 | 947 |
| **$p_{FWE}$** | 0.0444 | 0.0046* | 0.0072* | 0.0002* | 0.0104 |
| **Peak \|t\| (μ)** | 5.09 (-3.13) | 7.03 (-2.91) | 5.42 (+1.70) | 5.15 (+0.99) | 5.14 (+0.12) |
| **age (β, p)** | +0.024, 3.63e$^{-06}$ | +0.021, 1.83e$^{-05}$ | -0.014, 1.01e$^{-12}$ | -0.017, 1.48e$^{-14}$ | -0.004, .055 |
| **group (β, p)** | -0.177, .0001 | -0.235, 1.66e$^{-08}$ | +0.109, 2.16e$^{-08}$ | +0.084, 6.64e$^{-05}$ | +0.050, .004 |
| **group x age (β, p)** | -0.014, .039 | -0.019, .003 | +0.010, 4.38e$^{-05}$ | +0.010, .0006 | -0.0005, .848 |
| **sex (β, p)** | -0.025, .581 | +0.013, .748 | +0.085, 1.19e$^{-05}$ | +0.046, .027 | +0.060, .0006 |

**Table 8.** FWE-corrected displacement clusters, group x age contrast.

| **Hemi** | L | L | L | R |
|---|---|---|---|---|
| **ID** | c20 | c34 | c36 | c34 |
| **Vertices (n)** | 683 | 633 | 449 | 2,412 |

| | | | | |
|---|---|---|---|---|
| **$p_{FWE}$** | 0.0212 | 0.0228 | 0.0488 | 0.0002* |
| **Peak \|t\| (μ)** | 4.73 (-2.49) | 4.55 (+0.66) | 4.05 (+2.82) | 5.40 (+1.19) |
| **age (β, p)** | +0.020, $5.78e^{-07}$ | -0.018, $8.43e^{-14}$ | -0.020, .0007 | -0.022, $7.97e^{-21}$ |
| **group (β, p)** | -0.097, .0056 | +0.038, .074 | +0.027, .606 | +0.090, .0001 |
| **group x age (β, p)** | -0.022, $4.16e^{-05}$ | +0.011, .0003 | +0.026, .0011 | +0.013, $6.07e^{-05}$ |
| **sex (β, p)** | +0.020, .575 | +0.085, $5.48e\text{-}^{05}$ | -0.034, .516 | +0.014, .539 |

**Table 9.** Significant clusters from vertex-wise displacement analysis.

| | **age** | | | | **group** | | | | | **groupXage** | | | |
|---|---|---|---|---|---|---|---|---|---|---|---|---|---|
| **hemi** | lh | lh | rh | rh | lh | lh | lh | rh | rh | lh | lh | lh | rh |
| **cluster_id** | c1 | c22 | c27 | c97 | c3 | c12 | c30 | c1 | c24 | c20 | c34 | c36 | c34 |
| **n_vertices** | 4916 | 627 | 4467 | 634 | 408 | 752 | 736 | 2235 | 947 | 683 | 633 | 449 | 2412 |
| **cluster_p_perm** | $2e^{-04}$ | 0.015 | $2e^{-04}$ | 0.026 | 0.044 | 0.005 | 0.007 | $2e^{-04}$ | 0.01 | 0.021 | 0.023 | 0.049 | $2e^{-04}$ |
| **peak_abs_t** | 7.902 | 4.328 | 9.653 | 7.175 | 5.091 | 7.036 | 5.421 | 5.147 | 5.139 | 4.735 | 4.556 | 4.048 | 5.405 |
| **mean_t** | -0.205 | -2.981 | -0.399 | -0.033 | -3.133 | -2.908 | 1.701 | 0.995 | 0.120 | -2.494 | 0.663 | 2.828 | 1.188 |
| **n_subjects** | 440 | 440 | 440 | 440 | 440 | 440 | 440 | 440 | 440 | 440 | 440 | 440 | 440 |
| **n_22q** | 190 | 190 | 190 | 190 | 190 | 190 | 190 | 190 | 190 | 190 | 190 | 190 | 190 |
| **n_HC** | 250 | 250 | 250 | 250 | 250 | 250 | 250 | 250 | 250 | 250 | 250 | 250 | 250 |
| **n_sessions** | 874 | 874 | 874 | 874 | 874 | 874 | 874 | 874 | 874 | 874 | 874 | 874 | 874 |
| **n_sessions_22q** | 465 | 465 | 465 | 465 | 465 | 465 | 465 | 465 | 465 | 465 | 465 | 465 | 465 |
| **n_sessions_HC** | 409 | 409 | 409 | 409 | 409 | 409 | 409 | 409 | 409 | 409 | 409 | 409 | 409 |
| **age_min** | 5.107 | 5.107 | 5.107 | 5.107 | 5.107 | 5.107 | 5.107 | 5.107 | 5.107 | 5.107 | 5.107 | 5.107 | 5.107 |
| **age_max** | 34.819 | 34.819 | 34.819 | 34.819 | 34.819 | 34.819 | 34.819 | 34.819 | 34.819 | 34.819 | 34.819 | 34.819 | 34.819 |
| **age_mean** | 16.867 | 16.867 | 16.867 | 16.867 | 16.867 | 16.867 | 16.867 | 16.867 | 16.867 | 16.867 | 16.867 | 16.867 | 16.867 |
| **beta_group** | -0.026 | 0.024 | 0.043 | -0.016 | -0.177 | -0.235 | 0.109 | 0.084 | 0.050 | -0.097 | 0.038 | 0.027 | 0.090 |
| **t_group** | -2.370 | 0.474 | 4.319 | -3.024 | -3.873 | -5.644 | 5.598 | 3.989 | 2.869 | -2.767 | 1.785 | 0.515 | 3.803 |
| **p_group** | 0.018 | 0.6357 | 0.0000 | 0.0025 | 0.0001 | 0.0000 | 0.0000 | 0.0001 | 0.0041 | 0.0057 | 0.0743 | 0.6068 | 0.0001 |
| **beta_age** | -0.002 | -0.020 | -0.008 | -0.001 | 0.024 | 0.021 | -0.014 | -0.017 | -0.004 | 0.020 | -0.018 | -0.020 | -0.023 |
| **t_age** | -1.422 | -3.606 | -6.747 | -2.366 | 4.631 | 4.284 | -7.129 | -7.689 | -1.918 | 4.998 | -7.463 | -3.375 | -9.360 |
| **p_age** | 0.155 | 0.0003 | 0.0000 | 0.0180 | 0.0000 | 0.0000 | 0.0000 | 0.0000 | 0.0551 | 0.0000 | 0.0000 | 0.0007 | 0.0000 |
| **beta_interaction** | -0.001 | 0.023 | 0.006 | 0.000 | -0.014 | -0.019 | 0.011 | 0.010 | 0.000 | -0.022 | 0.012 | 0.026 | 0.013 |
| **t_interaction** | -0.370 | 3.046 | 3.818 | 0.594 | -2.055 | -2.931 | 4.086 | 3.407 | 0.192 | -4.099 | 3.583 | 3.268 | 4.010 |

| | | | | | | | | | | | | | |
|---|---|---|---|---|---|---|---|---|---|---|---|---|---|
| **p_interaction** | 0.712 | 0.0023 | 0.0001 | 0.5528 | 0.0399 | 0.0034 | 0.0000 | 0.0007 | 0.8480 | 0.0000 | 0.0003 | 0.0011 | 0.0001 |
| **beta_sex** | 0.034 | -0.029 | 0.010 | 0.010 | -0.025 | 0.013 | 0.085 | 0.046 | 0.060 | 0.020 | 0.085 | -0.034 | 0.014 |
| **p_sex** | 0.002 | 0.562 | 0.325 | 0.064 | 0.582 | 0.748 | 0.000 | 0.027 | 0.001 | 0.575 | 0.000 | 0.517 | 0.540 |
| **cohens_d_age5** | -0.122 | -0.363 | -0.178 | -0.311 | -0.022 | -0.016 | -0.085 | -0.137 | 0.192 | 0.334 | -0.337 | -0.389 | -0.217 |
| **cohens_d_age10** | -0.142 | -0.195 | 0.035 | -0.278 | -0.136 | -0.176 | 0.145 | 0.054 | 0.203 | 0.109 | -0.140 | -0.208 | 0.008 |
| **cohens_d_age15** | -0.162 | -0.026 | 0.248 | -0.245 | -0.251 | -0.335 | 0.374 | 0.246 | 0.214 | -0.116 | 0.058 | -0.027 | 0.233 |
| **cohens_d_age20** | -0.183 | 0.143 | 0.461 | -0.211 | -0.366 | -0.495 | 0.604 | 0.437 | 0.225 | -0.340 | 0.255 | 0.153 | 0.458 |
| **cohens_d_age25** | -0.203 | 0.311 | 0.674 | -0.178 | -0.481 | -0.654 | 0.833 | 0.629 | 0.235 | -0.565 | 0.453 | 0.334 | 0.683 |
| **cohens_d_age30** | -0.223 | 0.480 | 0.887 | -0.144 | -0.596 | -0.814 | 1.062 | 0.820 | 0.246 | -0.790 | 0.651 | 0.515 | 0.908 |
| **cohens_d_age35** | -0.244 | 0.649 | 1.100 | -0.111 | -0.711 | -0.974 | 1.292 | 1.012 | 0.257 | -1.015 | 0.848 | 0.695 | 1.133 |
| **sigma_u** | 0.037 | 0.205 | 0.050 | 0.029 | 0.236 | 0.096 | 0.135 | 0.124 | 0.093 | 0.110 | 0.079 | 0.209 | 0.163 |
| **sigma_e** | 0.150 | 0.648 | 0.123 | 0.061 | 0.558 | 0.590 | 0.197 | 0.235 | 0.212 | 0.482 | 0.283 | 0.695 | 0.236 |

#### 4 Claustrum Surface Displacement and Psychopathology

We further tested the clinical implications of surface displacement for 22q11DS carriers with prodromal psychotic symptoms (PPS+, N = 98), anxiety disorders (GAD, Separation Anxiety, Specific/Social Phobia; N = 104), or neurodevelopmental disorder (ADHD, N = 93). No clusters survived in subjects with psychotic symptoms or ADHD, in either hemisphere.

No clusters survived for lifetime psychotic symptoms or for ADHD, in either hemisphere, for either displacement or thickness. The psychotic symptoms result used the broadest available definition, so narrower definitions (schizophrenia/psychosis spectrum) would be expected to yield the same null. The concordance of the two geometrically independent measures indicates the absence of association is not specific to a single morphometric measure.

Anxiety was the only category showing any association. On displacement, carriers with a history of anxiety disorder showed a single FWE-significant age-modulated effect in the right hemisphere (c44, 552 vertices, $p$ = 0.046); no anxiety subtype (specific phobia, social phobia, GAD, separation anxiety) survived independently, and no main effect survived. This cluster fell entirely within the normative right age territory and almost entirely within the right group x age cluster (c34, 98% within), indicating it occupied surface already implicated in typical development and diagnostic divergence rather than an anxiety-specific region. Consistent with this, adding anxiety as a covariate to the c34 cluster-mean model left the group x age coefficient essentially unchanged, and the within-22q11DS anxiety x age coefficient was non-significant.

On thickness, anxiety yielded a left hemisphere main effect (c100, $p$ = 0.020) and a right hemisphere age-modulated effect (c69, $p$ = 0.015). Given a small number of marginal clusters, no subtype specificity, and spatial confounding of the displacement effect with the normative developmental territory, this association is interpreted as weak and exploratory.

#### 5 Claustrum-Working Memory Association Robustness

We evaluated three alternative explanations for the absence of claustrum-working memory association in 22q11DS carriers.

*Measurement range.* The absent association was not attributable to restricted variance or a floor effect in DSB. In 22q11DS carriers, DSB showed substantial individual variation (n = 360 sessions; mean = 5.7, SD = 2.3, range 0–14; only 1% at the floor, 14 distinct values), in a larger within-group sample than HC (174 vs 152 subjects). DSB also correlated with general cognitive ability within carriers (FSIQ, r = 0.34), confirming it as a valid individual difference measure. The carrier association estimates were near zero rather than imprecise, inconsistent with a floor effect or power explanation.

*Statistical equivalence.* To test whether the carrier association was positively equivalent to zero rather than merely non-significant, we applied a two-one-sided-tests (TOST) equivalence procedure with the control effect size as the equivalence bound. In all five significant DSB x age clusters the carrier coupling was statistically equivalent to zero (TOST $p \leq 0.004$), and each carrier 95% confidence interval excluded the control estimate.

*Participant age.* Carriers were modestly older than controls (mean 17.5 vs 15.7 years; Welch t, $p < 0.001$), and the HC association was age-graded (stronger in childhood). To ensure the non-association was not an age artifact, we compared the association between groups at matched age using a full DSB x group x age model (group difference evaluated at the centered age) and in an age-balanced subsample (inverse probability weighting on age). The group difference remained significant at matched age in four of five clusters ($p$ = 0.005–0.038) and showed the same direction in the age-balanced subsample, attenuating only slightly relative to the age-unadjusted estimate. The decoupling therefore does not depend on the group age difference.

*Antipsychotic exposure.* Because antipsychotic medication can alter brain structure, we tested whether it accounted for the non-association. Antipsychotic use was rare and confined to carriers (5% of 22q11DS sessions, 22/419). Repeating the decoupling analysis after excluding all antipsychotic-exposed sessions left the result unchanged; the group difference in coupling remained significant in four of five clusters ($p$ = 0.004–0.029), indicating the decoupling is not attributable to medication.
